# SARS-CoV-2 Variant Identification Using a Genome Tiling Array and Genotyping Probes

**DOI:** 10.1101/2021.05.11.443659

**Authors:** Ryota Shimada, Emily N. Alden, Kendall Hoff, Xun Ding, Jiayi Sun, Adam M. Halasz, Wei Zhou, Jeremy S. Edwards

## Abstract

With over three million deaths worldwide attributed to the respiratory disease COVID-19 caused by the novel coronavirus SARS-CoV-2, it is essential that continued efforts be made to track the evolution and spread of the virus globally. We previously presented a rapid and cost-effective method to sequence the entire SARS-CoV-2 genome with 95% coverage and 99.9% accuracy. This method is advantageous for identifying and tracking variants in the SARS-CoV-2 genome when compared to traditional short read sequencing methods which can be time consuming and costly. Herein we present the addition of genotyping probes to our DNA chip which target known SARS-CoV-2 variants. The incorporation of the genotyping probe sets along with the advent of a moving average filter have improved our sequencing coverage and accuracy of the SARS-CoV-2 genome.

## INTRODUCTION

The novel Coronavirus, SARS-CoV-2, is attributed to over one 141 million cases of the respiratory disease COVID-19 and has resulted in over 3 million deaths globally since its first report in December of 2019^1^. SARS-CoV-2 is a positive sense, single stranded RNA virus prone to mutations in part due to its viral polymerase lacking error correction capabilities, resulting in poor replication fidelity^2,3,4^. Additionally, the 3’ to 5’ exoribonuclease activity has been attributed to moderate mutations in SARS-CoV-2 which affect transmission, receptor interactions, and host infectivity^5,6,7^. With the help of whole genome sequencing many variants in the viral genome have been detected, several of which may be associated with an increased risk of mortality^8,9,10^.

In our previous work we introduced a SARS-CoV-2 whole genome resequencing approach utilizing a DNA array chip that can sequence accurately, at single base-pair resolution, the complete SARS-CoV-2 genome^11^. Eight SARS-CoV-2 clinical samples from COVID-19 patients in Wyoming were sequenced and yielded a sequencing accuracy and coverage of 99.86%-99.92% and 98.07%-99.54% respectively. Here we introduce a DNA chip that includes genome tiling features, as previously described, plus specific probe sets to genotype known SARS-CoV-2 variants. The data from the additional probes sets allows for a confirmation measurement of variant calls made by the tiling array probes sets. In addition, we have further improved the variant identification accuracy via a moving-average based filtering algorithm which identifies regions of low-quality data and determines the reliability of the variant calls. These additions to our previous work have resulted in a notable improvement of sequencing accuracy and coverage of 99.92%-99.99% and 98.37%-99.59% respectively.

## Methods

### Sample Preparation and Hybridization

Samples were prepared as described in our previous work^11^. Briefly, in order to test the robustness and improved sequencing accuracy of the DNA chip by including specific genotyping probes in addition to the tiling array features, we sequenced eight clinical samples from COVID-19 positive patients. The eight clinical samples were acquired from the Wyoming Public Health Laboratories through collaboration with the UNM Department of Pediatrics as described in Hoff *et al*. We will maintain the naming scheme of our previous work and refer to these samples as WY24, WY26, WY32, WY36, WY41, WY44, WY59, and WY64.

The samples re-amplified using the ARTIC protocol^12^ using biotinylated dUTP in order to prepare the sample for sequencing using the DNA chip equipped with both the tiling DNA array and genotyping probe sets. Fragmented samples were hybridized to the DNA chip overnight prior to Cy3-Streptavidin staining. The chips were then scanned using a custom-built confocal scanner in the Cy3 channel.

### DNA Chip Design

The tiling array is identical to the DNA chip design described previously^11,13^. In short, each position in the genome (on the sense and antisense strand) is represented on the chip using a group of features composed of 25-mer oligonucleotide probes. Each feature in the probe sets is represented by one of the four bases at the 13th position, flanked on either side with the reference sequence. Thus, each position in the SARS-CoV-2 genome is represented by 2 probe sets, one for the sense strand and one for the antisense strand on the tiling DNA array. Note, SARS-CoV-2 is a ssRNA virus, but the antisense strand is synthesized during the ARTIC PCR amplification.

In addition to tiling array probes for every position of the SARS-CoV-2 genome, we incorporated probes specific to sites of known variants and genomic regions of interest. These positions were identified by analyzing all SARS-CoV-2 full genome sequences in the GISAID database as of May 2020 utilizing the cleaning and filtering functions designed by Lanfear^14,15^. Similar to the tiling array probes, each variant selected for genotyping was synthesized on the chip with probe sets for both the sense and antisense strands.

### Base or Variant calling method for genome tiling array and genotyping probe sets

Using the maximum likelihood method developed by Hoff 2021, the raw data from the genome tiling array was used to construct the maximum likelihood (ML) model variant and base calling and assigning the Q score each base call (either variant or reference base). The difference and differential were calculated for the genotyping probe sets. The difference and differential were calculated as described by Hoff et al; briefly, the difference is the highest intensity feature - lowest intensity feature, and the differential is the highest intensity feature - second highest intensity feature normalized by the difference. The Q score for the base or variant calls from the genotyping features was calculated using the ML model constructed from the tiling array data.

Accuracy and coverage calculations of the genome tiling array are used as the baseline for comparison purposes throughout our work. Accuracy calculations account for the number of correct reference calls, correctly identified true variant calls, and incorrect calls divided by the total number of high-quality calls where the Q score was greater than the Q-threshold (Qth) of 20. Coverage is calculated as the total number of calls (both reference base or variant calls) divided by the size of the SARS-CoV-2 reference genome.

### Incorporation of the Genotyping Probe-set

To improve overall accuracy and coverage we devised a method to incorporate high quality calls made by the genotyping probe-set, replacing low quality calls from the genome tiling array, while retaining the high-quality calls of the genome tiling array. The replacement strategy looks to improve the calls made by the genome tiling array in two separate iterations which look at the ‘correctness’ of the call relative to the reference and the overall Q score of each individual read. Specifically, the first iteration of replacement removes variant calls made by the tiling array that are not confirmed by genotype probes, where the genotyping Q score is higher. The second iteration of replacement serves to improve the overall quality of the entire sequencing data set by incorporating the Q score for the genotyping probes when the calls agree. The following sections will provide details on the utilization of the genotyping probes along with the tiling array features.

#### First Iteration of the Replacement and Incorporation of the Genotyping Probe-set

For all variant calls from the tiling array with a Q score less than 50, the base call for the corresponding position from the genotyping array is examined. If the Q score for the genotyping probe does not confirm the variant call and has a higher Q score, the variant call is removed, and a reference base is called. Additionally, during this set, if the tiling array does not call a base because the Q score is below 20, a reference or variant call is made based on the genotyping probes if the genotyping Q score is greater than 20.

#### Second Iteration of the Replacement and Incorporation of the Genotyping Probe-set

The second iteration will replace additional calls that have higher Q-scores in the genotyping probes than in the genome tiling array. This additional step improves the reliability of the previous reference calls and increases the total number of reads being used. For instance, when both the genome tiling array and the genotyping probe set make the same reference base call, the read with the higher Q score is selected. Additionally, when the genotyping probe set confirms a variant call made and has a higher Q score than the genome tiling array, then the data with the higher Q score is retained.

### Q score Moving Average Filter

We developed a simple tool in R to calculate the individual moving average of Q scores (MAQs) centered around each position in the SARS-CoV-2 genome. The MAQ was calculated for each genome position considering the twelve flanking bases on either side of the position, 25 bases total which is the length of the oligonucleotides on the chips. The calculated MAQ value for every position in the genome was added as metadata for every base call.

Using the MAQ, we developed a simple moving average filter (MA-filter) as an additional basecalling criterion to improve the reliability and confidence of the base calls made by the DNA chip. First, variant calls with a Q score above 30 were not analyzed by the MA-filter. However, variant calls with a Q score below 30 were subject to the MA-filter. We then filtered out and removed variant calls with a Q score below 30 if the MAQ was two standard deviations below the average of the Q scores across the entire genome. The base calls were then categorized as “Non-calls” and the data removed from the final analysis. The position, Q score, and MA of all reads identified as for the MA-filter-group were recorded.

The accuracy and coverage of the post-MA-filtered data was calculated as described in previous methods. The results were graphically compiled using ggplot2 package version 3.3.2 in R^16^.

### Accuracy analysis of samples

True variants were separately identified by Illumina short-read sequencing and used to determine the accuracy of the variant calls made by the combined genome tiling array and genotyping probe sets^11^. We then analyzes the accuracy of the results from the chip based sequencing using tiling array features and genotyping probes on four sets of genome positions; all sites in the SARS-CoV-2 genome, the 42 positions of known SARS-CoV-2 variants from the Wyoming clinical samples in the GISAID database as of May 2020, all US SARS-CoV-2 variants from the GISAID database as of May 2020 (US Var, 5055 positions) and confirmed US SARS-CoV-2 variants which have been identified in five or more samples as of May 2020 (USgt5, 1044 positions). The post replacement, MA-filtered sequencing data was assessed for basecalling accuracy at the positions described. We systematically assessed each known variant location and verified whether-or-not the genome tiling array and the genotyping probe set made the correct variant calls at these positions with a sufficient Q score. Variant calls that did not match the short-read sequencing true variant results were identified for further investigation. The base calls identified included reference calls made at true variant positions, incorrect variant calls at true variant positions, “Non-call” reads either due to low Q, or “Non-call” reads filtered out by the MA-filter. True variants were only correctly identified if the base call matched the variant base call of the Illumina short-read sequencing. A categorical Venn diagram of the variant calls was made using the VennDiagram package in R, version 1.6.20^17^.

## Results and Discussion

### Incorporation of genotyping probe sets and genome tiling array probe set data

Complementing the data from the genome tiling array probe sets with the genotyping probe sets increased the accuracy of the base calls by three different mechanisms. First, when the variant is in a region that is identical to the reference sequence except for a single nucleotide variant, the genotyping probes simply provide additional measurements of a base. Secondly, the variant may reside in a region with additional variants, so therefore, the surrounding ‘reference’ sequence of the tiling array is different from the surrounding sequence on the genotyping array. Finally, the genotyping probes can assay more complex variants, such as indels. In the clinical samples we analyzed all improvements in the accuracy were due to additional redundant measurements because. The redundant measurements likely improved the base calling accuracy due to issues with certain probes sets and the probe sets location on the chip. For example, nearby extremely bright features can lead to slightly erroneous intensity measurements, which likely leads to some incorrect variant calls.

Therefore, to improve the base and variant calling accuracy, we incorporated the base calls from the genotyping probes with the data from the genome tiling array probes in two iterations. The first iteration replaces non-reference calls and non-calls made by the genome tiling array with reference calls made by the genotyping probe set when the data quality from the genotyping probe sets is of higher quality. Table 1 illustrates the results after the first iteration of replacements. Overall, for the WY64 samples, the number of variant calls decreased by 2 when incorporating the genotyping probe set data. In the WY64 sample, we made 23 variant calls when using only the genome tiling array, and after the first iteration of replacements we identified 2 base calls that should be a reference call rather than a variant. Also, the coverage of the WY64 sample increased from 99.538% to 99.554% by replacing non-calls with a genotyping probe call. Finally, with the second iteration of replacement additional base calls that have been determined by the genotyping probe sets to have a higher Q score where adjusted. This additional step improves the reliability of the previous reference calls and expands the genome coverage to 99.6180% respectively (Table 1).

**Table 1.**
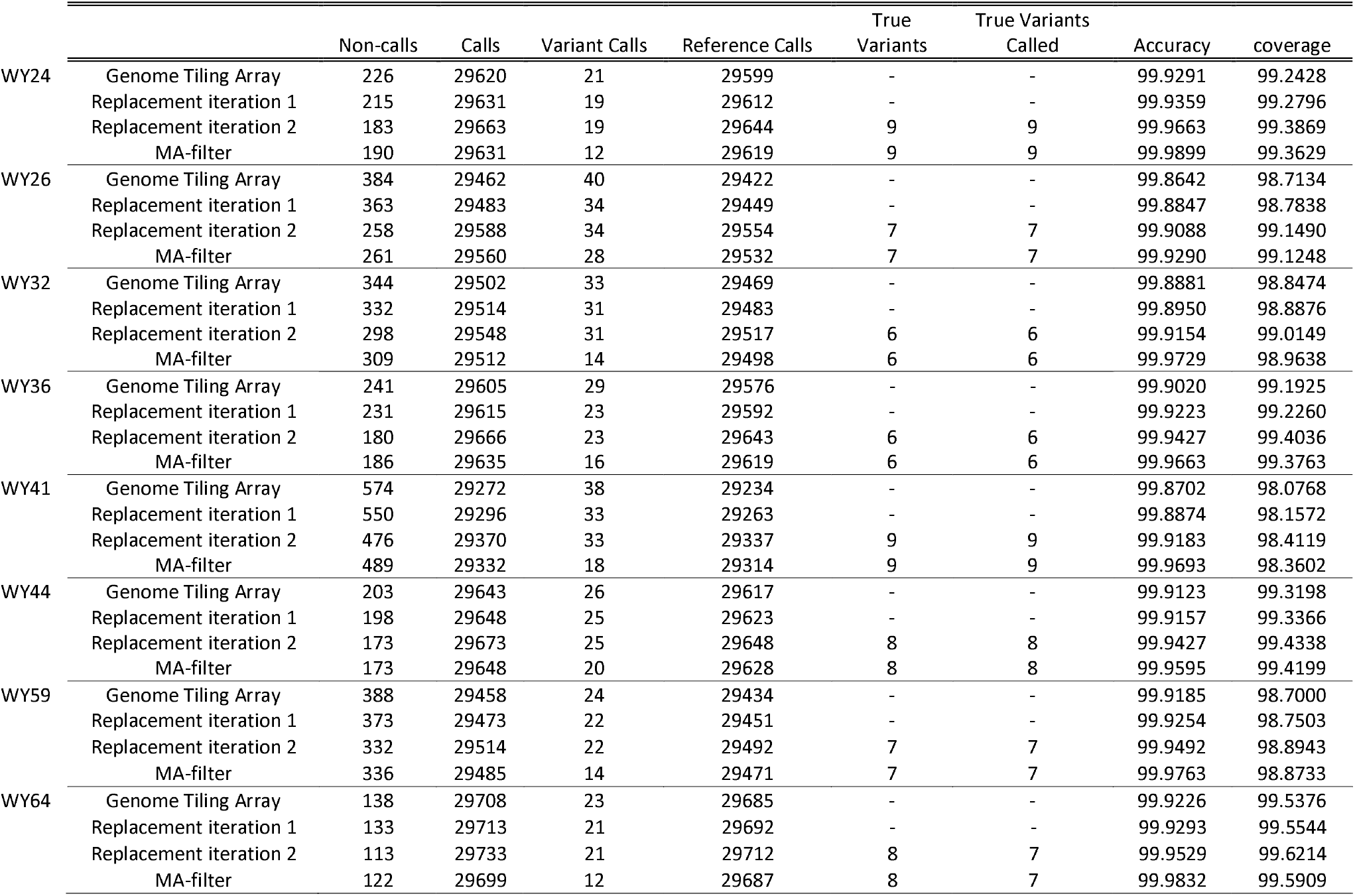
Incremental accuracy improvements of base calls made by the DNA tiling array. Results and breakdown of call accuracy, coverage, and viability of reads made with the genome tiling array and improvements made through the replacement iteration with the genotyping probe set data and MA-filter. Non-calls refer to calls where Q-score is less than Qth = 20 and/or calls that are filtered out by MA-filter. Calls refer to base calls where Q-score is greater than Qth and used for sequencing calls. Variant calls refer to non-reference base-calls. Reference calls are those made relative to the reference genome. True variants called are correct calls made in reference to known true variants. Accuracy and coverage are both percentage values.

|  |  | Non-calls | Calls | Variant Calls | Reference Calls | True<br>Variants | True Variants<br>Called | Accuracy | coverage |
| --- | --- | --- | --- | --- | --- | --- | --- | --- | --- |
| WY24 | Genome Tiling Array | 226 | 29620 | 21 | 29599 | - | - | 99.9291 | 99.2428 |
|  | Replacement iteration 1 | 215 | 29631 | 19 | 29612 | - | - | 99.9359 | 99.2796 |
|  | Replacement iteration 2 | 183 | 29663 | 19 | 29644 | 9 | 9 | 99.9663 | 99.3869 |
|  | MA-filter | 190 | 29631 | 12 | 29619 | 9 | 9 | 99.9899 | 99.3629 |
| WY26 | Genome Tiling Array | 384 | 29462 | 40 | 29422 | - | - | 99.8642 | 98.7134 |
|  | Replacement iteration 1 | 363 | 29483 | 34 | 29449 | - | - | 99.8847 | 98.7838 |
|  | Replacement iteration 2 | 258 | 29588 | 34 | 29554 | 7 | 7 | 99.9088 | 99.1490 |
|  | MA-filter | 261 | 29560 | 28 | 29532 | 7 | 7 | 99.9290 | 99.1248 |
| WY32 | Genome Tiling Array | 344 | 29502 | 33 | 29469 | - | - | 99.8881 | 98.8474 |
|  | Replacement iteration 1 | 332 | 29514 | 31 | 29483 | - | - | 99.8950 | 98.8876 |
|  | Replacement iteration 2 | 298 | 29548 | 31 | 29517 | 6 | 6 | 99.9154 | 99.0149 |
|  | MA-filter | 309 | 29512 | 14 | 29498 | 6 | 6 | 99.9729 | 98.9638 |
| WY36 | Genome Tiling Array | 241 | 29605 | 29 | 29576 | - | - | 99.9020 | 99.1925 |
|  | Replacement iteration 1 | 231 | 29615 | 23 | 29592 | - | - | 99.9223 | 99.2260 |
|  | Replacement iteration 2 | 180 | 29666 | 23 | 29643 | 6 | 6 | 99.9427 | 99.4036 |
|  | MA-filter | 186 | 29635 | 16 | 29619 | 6 | 6 | 99.9663 | 99.3763 |
| WY41 | Genome Tiling Array | 574 | 29272 | 38 | 29234 | - | - | 99.8702 | 98.0768 |
|  | Replacement iteration 1 | 550 | 29296 | 33 | 29263 | - | - | 99.8874 | 98.1572 |
|  | Replacement iteration 2 | 476 | 29370 | 33 | 29337 | 9 | 9 | 99.9183 | 98.4119 |
|  | MA-filter | 489 | 29332 | 18 | 29314 | 9 | 9 | 99.9693 | 98.3602 |
| WY44 | Genome Tiling Array | 203 | 29643 | 26 | 29617 | - | - | 99.9123 | 99.3198 |
|  | Replacement iteration 1 | 198 | 29648 | 25 | 29623 | - | - | 99.9157 | 99.3366 |
|  | Replacement iteration 2 | 173 | 29673 | 25 | 29648 | 8 | 8 | 99.9427 | 99.4338 |
|  | MA-filter | 173 | 29648 | 20 | 29628 | 8 | 8 | 99.9595 | 99.4199 |
| WY59 | Genome Tiling Array | 388 | 29458 | 24 | 29434 | - | - | 99.9185 | 98.7000 |
|  | Replacement iteration 1 | 373 | 29473 | 22 | 29451 | - | - | 99.9254 | 98.7503 |
|  | Replacement iteration 2 | 332 | 29514 | 22 | 29492 | 7 | 7 | 99.9492 | 98.8943 |
|  | MA-filter | 336 | 29485 | 14 | 29471 | 7 | 7 | 99.9763 | 98.8733 |
| WY64 | Genome Tiling Array | 138 | 29708 | 23 | 29685 | - | - | 99.9226 | 99.5376 |
|  | Replacement iteration 1 | 133 | 29713 | 21 | 29692 | - | - | 99.9293 | 99.5544 |
|  | Replacement iteration 2 | 113 | 29733 | 21 | 29712 | 8 | 7 | 99.9529 | 99.6214 |
|  | MA-filter | 122 | 29699 | 12 | 29687 | 8 | 7 | 99.9832 | 99.5909 |

### Moving average filter to identify unreliable variant calls

A moving average (MA) filter was developed as a means to objectively identify base calls that might be less reliable. This method was developed because we previously observed a high number of variant calls of low quality in certain regions of the genome. Therefore, a region of low MA Q score indicates that the data is less reliable. After the incorporation of the genotyping probes, we made 21 variant calls for WY64, and 9 of these variant calls had a Q score between 20 and 30 and were flagged for further analysis. The average Q score across the entire genome in sample WY64 was 40.7004 (S.D. 2.2559), and we therefore set the MA Q score cutoff at two standard deviations below the mean, 36.1886. All 9 reads flagged for further analysis had a MA Q score lower than the cutoff and thus were removed from the analysis. Ultimately, in the WY64 sample we identified 113 non-calls with a Q score less than 20 prior to the MA-filter. The number of non-calls increased to 122 after the 9 lower quality calls were removed and the number of variant calls were decreased to 12. The categorical breakdown of the “called” reads as a function of Q-score and position is depicted as a scatter plot in Figure 1.

**Figure 1.**
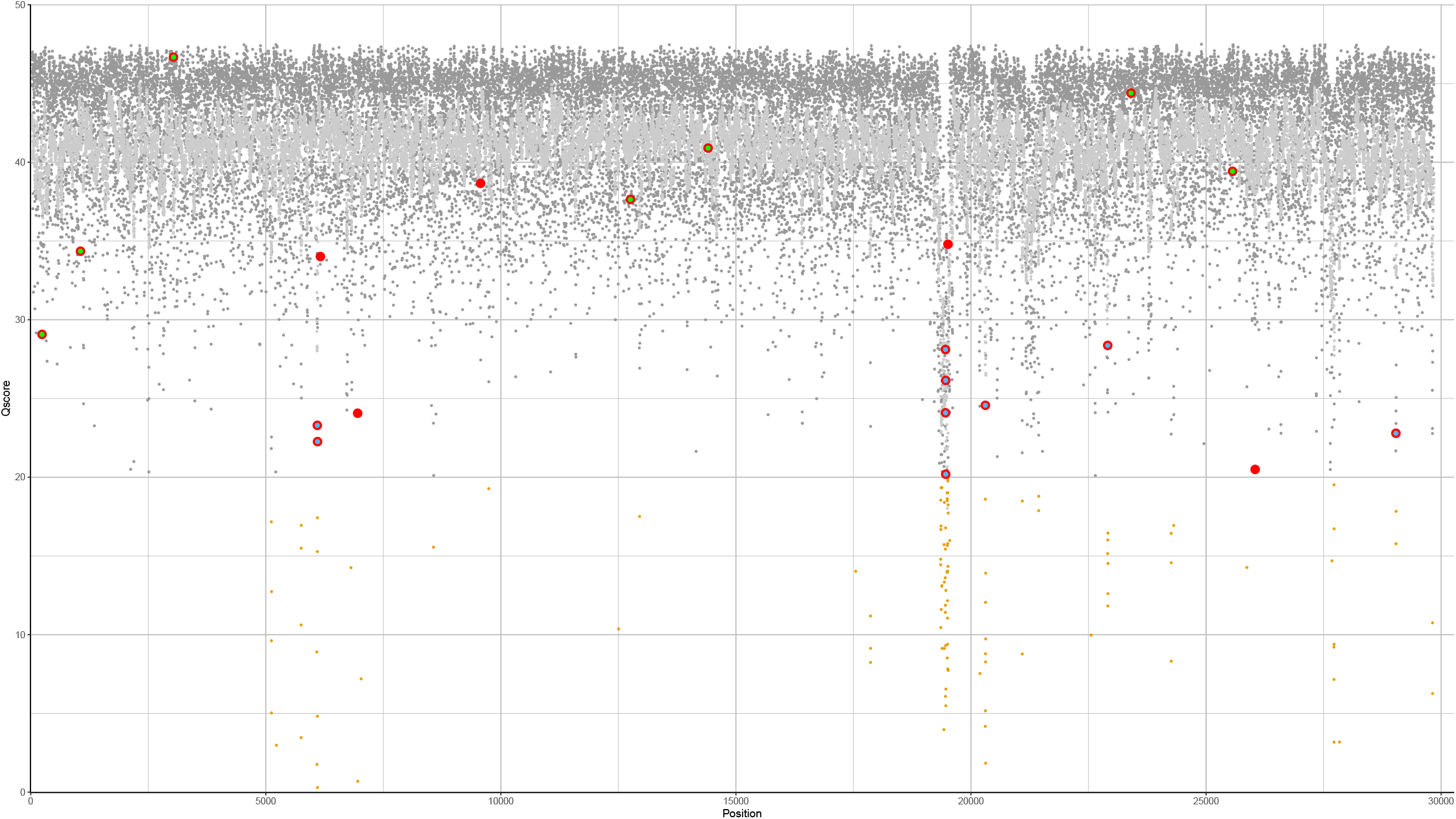
Scatter plot, WY64 Q score and MAQ with variant call breakdown after MA-filter. A scatter plot generated in R using ggplot2 displaying all reads made by the full genome tiling array reads with incorporation of genotyping probe-set data and MA-filter on sample WY64 between position 26 and 29834. Every call was assessed with a combination of base call and Q score. High quality reads with Q > Qth (20) are used to make calls. Dark gray circles represent base positions where Q > Qth and make a ‘Reference Call’. All calculated MAQ are overlaid as light gray. Brown dots are reads that have a low Q score where Q < Qth, are categorically ‘Non-calls’, and excluded to make base-calls. Q scores of ‘Variant Calls’ are identified as larger red circles, where the final base call made by DNA chip after replacement and MA-filter is not reference and the Q > Qth. Within ‘Variant Calls’, a blue overlap indicates calls that are removed by the MA-filter, which takes all variant calls with a Q score between 20 and 30 and removes any with a MAQ lower than the MA-threshold. Within ‘Variant Calls’, a green overlap indicates true variant calls identified and verified with short-read sequencing data.

### Validation of variant calls

To verify the variant calls, we compared the resequencing results from the full-genome tiling array and genotyping probe sets after applying the MA-filter, to both the reference genome and to the variant calls from the Illumina sequencing results. For the WY64 sample, we confirmed whether a variant call made by the resequencing chip correctly detects a variant (matches Illumina sequencing results), makes an erroneous variant call (Illumina data calls a reference base), or misses a variant that Illumina identified. In this analysis we assumed that the Illumina data is 100% accurate at identifying viral variants.

For the WY64 sample, we missed one true variant at position 24,453 in the genome (Fig. 2). This variant was initially detected by the genome tiling array with a relatively low Q score of 24.8; however, the genotyping probe sets for this position called a reference base with a relatively low, but higher Q score of 25.3. Therefore, we ultimately called a reference base at this position. To further investigate the base at this position, we looked at the variant call file that was produced during the analysis of the Illumina short read data. The Illumina short read alignments at this position indicated that 251 mapped reads covered this location in the SARS-CoV-2 genome. Of these 251 reads, 127 contained the reference base at this position, while 124 reads contained the variant base at this position. Therefore, we suspect that this clinical sample contains quasispecies; some viruses have the 24,453 variant while others represent the reference genome. The existence of quasispecies may explain the relatively low Q score for this position, which is well below the average Q score seen across all positions in the genome (Q>40).

**Figure 2.**
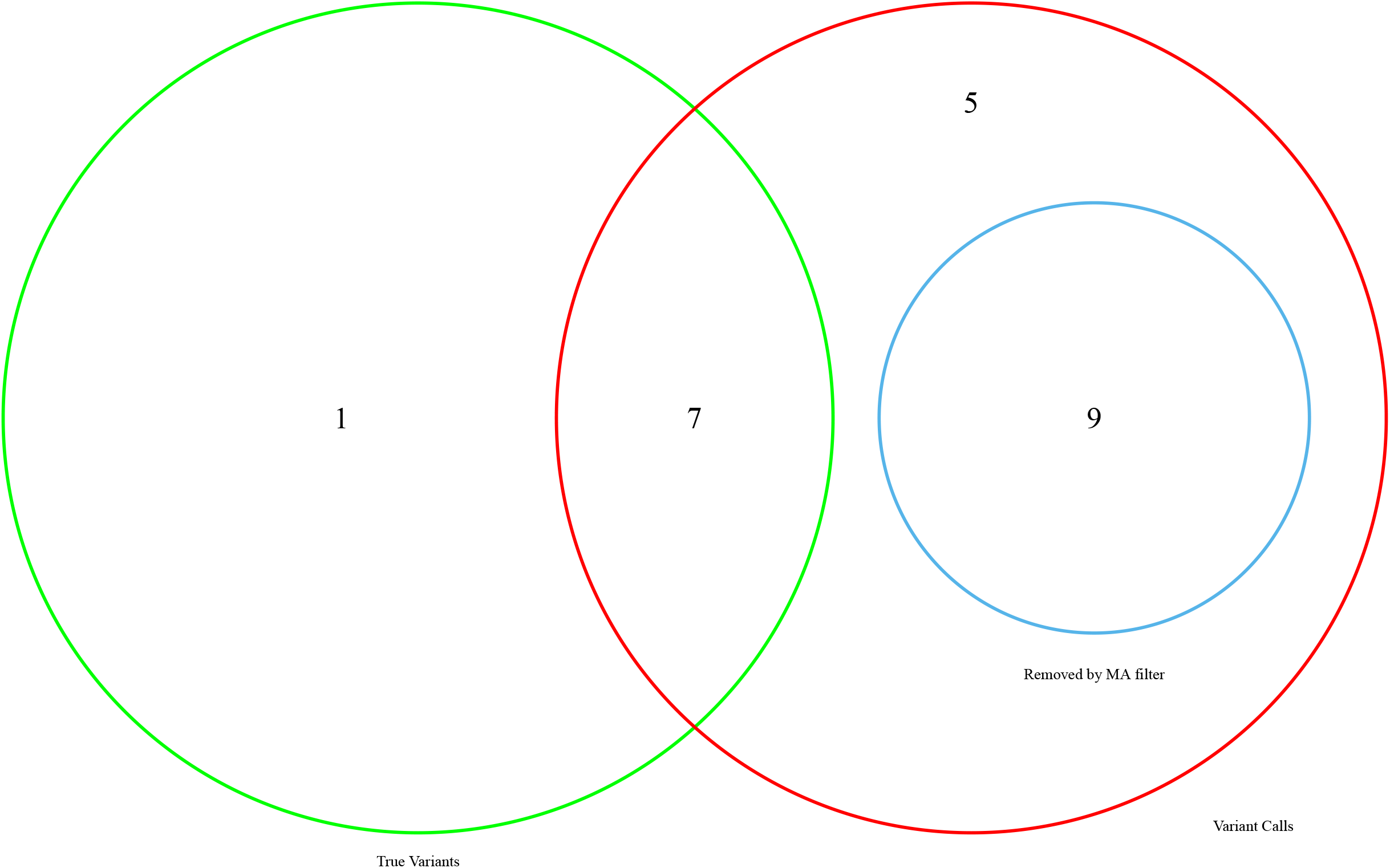
Venn Diagram, WY64 Variant Calls after MA-filter. A Venn diagram depicting the categorical breakdown of calls made by the DNA chip on sample WY64. Variant Calls made by the DNA chip are contained within the red circle. Any circle or number outside of red implies that the call is a reference call. True variants are color coded in green and are confirmed by short-read sequencing data. The blue circle indicates variant calls that are filtered out by the MA-filter, converted to non-calls, and omitted from making base calls. Overlap between green and red circles indicates true variants correctly called by the DNA chip and verified through short-read sequencing data. Overlap between green and blue circles indicate an improperly removed true variant by the MA-filter where the read is situated in a local region of low MAs indicated by a low MAQ below the MAQ threshold. Ideally both green and blue circles should be contained within the red circle without overlapping each other, implying that all true variants and reads filtered out by the MA-filter were correctly identified as variant calls.

While we speculate that our inability to correctly call the variant at position 24,453 in the WY64 sample is due to the presence of a quasispecies of virus, further investigation is needed. Viral quasispecies arise due to the high mutation rates of the viral genome and the change of relative frequency of the variants in the population^18^. Quasispecies have been identified in the novel SARS-CoV-2 virus and their associated mutations have been shown to have a moderate to high impact on viral gene expression^19^. We hypothesize that simultaneous high Q scores in genome tiling array and genotyping probes at a single base may be an indication of the existence of quasispecies, and our data analysis pipeline can be extended in the future to quantify the presence of quasispecies.

### Genotyping at known variants site

To examine how effective our approach is at genotyping known variant sites, we analyzed the accuracy of the results at the known variant sites within all Wyoming samples, all United States samples, and also all variants sites that appear more than five times in the United States samples. The definition of known sites is based on data within the GISAID database as of August 2020.

Using the maximum likelihood method, replacement methods, and removal of low-quality reads through the MA-filter, we analyzed 8 SARS-CoV-2 genomes from clinical samples at the sites of known variants. Overall, seven of the eight SARS-CoV-2 samples had 100% accuracy at the 42 identified WY variant positions after the replacement iterations and MA-filter. In sample 64, we replaced a single correct variant call with an erroneous reference call, leading to an accuracy of 97.62% at the 42 Identified WY variant positions. However, we suspect this error may be due to the presence of a quasispecies. Accuracy at the USgt5 variant positions ranged between 99.70% to 100% (WY32 and WY36 respectively) and 99.68%-99.94% accuracy at all US-variant positions (WY32 and WY24, respectively) (Table 2).

**Table 2.** Accuracy improvements of genotyping at known variant locations. Results of progressive improvements by the replacement and MA-filter methods at specific variant size, organized by sample. WY var examines 42 base positions of known variants within samples gathered and sequenced in WY. US-gt5 are 1044 sites of variants sequenced, detected and confirmed in the US that have occurred in more than 5 samples. US var are sites of all 5055 variants found in US samples as of August 2020 on the GISAID database. Accuracy calculations are confined exclusively to the variant locations

|  |  | Genome<br>Tiling Array | Replacement<br>Iteration 1 | Replacement<br>Iteration 2 | MA-filter |
| --- | --- | --- | --- | --- | --- |
| WY24 | WY var | 100.00% | 100.00% | 100.00% | 100.00% |
|  | US-gt5 | 99.61% | 99.90% | 99.90% | 99.90% |
|  | US var | 99.86% | 99.94% | 99.94% | 99.94% |
| WY26 | WY var | 100.00% | 100.00% | 100.00% | 100.00% |
|  | US-gt5 | 99.51% | 99.80% | 99.80% | 99.80% |
|  | US var | 99.56% | 99.76% | 99.76% | 99.78% |
| WY32 | WY var | 100.00% | 100.00% | 100.00% | 100.00% |
|  | US-gt5 | 99.51% | 99.51% | 99.51% | 99.70% |
|  | US var | 99.58% | 99.62% | 99.62% | 99.68% |
| WY36 | WY var | 100.00% | 100.00% | 100.00% | 100.00% |
|  | US-gt5 | 99.90% | 99.90% | 99.90% | 100.00% |
|  | US var | 99.78% | 99.84% | 99.84% | 99.88% |
| WY41 | WY var | 100.00% | 100.00% | 100.00% | 100.00% |
|  | US-gt5 | 99.80% | 99.80% | 99.80% | 99.80% |
|  | US var | 99.58% | 99.70% | 99.70% | 99.72% |
| WY44 | WY var | 100.00% | 100.00% | 100.00% | 100.00% |
|  | US-gt5 | 99.61% | 99.61% | 99.61% | 99.80% |
|  | US var | 99.76% | 99.80% | 99.80% | 99.88% |
| WY59 | WY var | 100.00% | 100.00% | 100.00% | 100.00% |
|  | US-gt5 | 99.80% | 99.80% | 99.80% | 99.90% |
|  | US var | 99.70% | 99.76% | 99.76% | 99.78% |
| WY64 | WY var | 100.00% | 97.62% | 97.62% | 97.62% |
|  | US-gt5 | 100.00% | 99.90% | 99.90% | 99.90% |
|  | US var | 99.90% | 99.92% | 99.92% | 99.92% |

### Conclusion

Genomic data of the SARS-CoV-2 virus is essential for combatting this global pandemic. These data have been used to create healthcare policy, create hospital protocols, for allocation of finite resources on the front lines, and the development of multiple vaccines. Our technology addresses the need for continued monitoring of viral evolution and the spread of known viral variants which is imperative in combating further outbreaks of the virus. Through our previous work on a genome tiling array for SARS-CoV-2, Hoff 2021, and the subsequent improvements by inclusion of the genotyping probe sets outlined in this paper, we provide a highly accurate, reliable, rapid, and cost-effective method for surveillance of the rapidly-mutating SARS-CoV-2 viral strains. By incorporating probes to detect known variants and a moving average filter to address low quality regions, we have sequenced eight clinical samples and improved the accuracy of our previous work from 99.86% to 99.98% and with improved coverage from 99.18% to 99.59%.

## Supporting information

Supplemental Figures and Legends for all Clinical Samples

## Acknowledgements

N.H. and D.D. provided access to the clinical samples. This research was partially supported by the UNM Comprehensive Cancer Center Support Grant NCI (P30CA118100) and an Institutional Development Award (IDeA) from the National Institute of General Medical Sciences of the National Institutes of Health (P20GM103451).

We also gratefully acknowledge the following authors found in Supplemental Table 1 from the originating laboratories responsible for obtaining the specimens and the submitting laboratories where genetic sequence data were generated and shared via the GISAID Initiative, on which this research is based.

## Author Contributions

#R.S. and E.A. contributed equally to this work.

### Author Contributions

K.H. and X.D. conducted the experiments. J.S. designed and constructed the genome arrays. R.S., E.N.A, and A.M.H. developed the base calling methods. J.S.E. and W.Z. conceived of and directed the study. All authors drafted the manuscript.

### Notes

The authors declare the following competing financial interest(s): The Centrillion affiliated authors are employees and the company may commercialize the work described herein.

## References

1. COVID-19 Map https://coronavirus.jhu.edu/map.html (accessed Feb 19, 2021).

2. Drew, TW.;. The Emergence and Evolution of Swine Viral Diseases: To What Extend Have Husbandry Systems and Global Trade Contributed to Their Distribution and Diversity? 2011, 30 (1), 95.

3. Severe acute respiratory syndrome coronavirus-2 (SARS-CoV-2): a global pandemic and treatment strategies https://www.ncbi.nlm.nih.gov/pmc/articles/PMC7286265/ (accessed Feb 19, 2021).

4. Domingo, E.; Martin, V.; Perales, C.; Grande-Pérez, A.; García-Arriaza, J.; Arias, A. Viruses as Quasispecies: Biological Implications. Curr Top Microbiol Immunol 2006, 299, 51–82. https://doi.org/10.1007/3-540-26397-7_3.

5. Minskaia, E.; Hertzig, T.; Gorbalenya, A. E.; Campanacci, V.; Cambillau, C.; Canard, B.; Ziebuhr, J. Discovery of an RNA Virus 3’→5’ Exoribonuclease That Is Critically Involved in Coronavirus RNA Synthesis. PNAS 2006, 103 (13), 5108–5113. https://doi.org/10.1073/pnas.0508200103.

6. Wan, Y. Receptor Recognition by the Novel Coronavirus from Wuhan: an Analysis Based on Decade-Long Structural Studies of SARS Coronavirus | Journal of Virology https://jvi.asm.org/content/94/7/e00127-20 (accessed Apr 21, 2021).

7. Ogando, N. S.; Zevenhoven-Dobbe, J. C.; Meer, Y. van der; Bredenbeek, P. J.; Posthuma, C. C.; Snijder, E. J. The Enzymatic Activity of the Nsp14 Exoribonuclease Is Critical for Replication of MERS-CoV and SARS-CoV-2. Journal of Virology 2020, 94 (23). https://doi.org/10.1128/JVI.01246-20.

8. Davies, N. G.; Jarvis, C. I.; Edmunds, W. J.; Jewell, N. P.; Diaz-Ordaz, K.; Keogh, R. H. Increased Mortality in Community-Tested Cases of SARS-CoV-2 Lineage B.1.1.7. Nature 2021, 1–5. https://doi.org/10.1038/s41586-021-03426-1.

9. Roussel, Y.; Giraud-Gatineau, A.; Jimeno, M.-T.; Rolain, J.-M.; Zandotti, C.; Colson, P.; Raoult, D. SARS-CoV-2: Fear versus Data. International Journal of Antimicrobial Agents 2020, 55 (5), 105947. https://doi.org/10.1016/j.ijantimicag.2020.105947.

10. Hauser, A.; Counotte, M. J.; Margossian, C. C.; Konstantinoudis, G.; Low, N.; Althaus, C. L.; Riou, J. Estimation of SARS-CoV-2 Mortality during the Early Stages of an Epidemic: A Modeling Study in Hubei, China, and Six Regions in Europe. PLOS Medicine 2020, 17 (7), e1003189. https://doi.org/10.1371/journal.pmed.1003189.

11. Hoff, K.; Ding, X.; Carter, L.; Duque, J.; Lin, J.-Y.; Dung, S.; Singh, P.; Sun, J.; Crnogorac, F.; Swaminathan, R.; Alden, E. N.; Zhu, X.; Shimada, R.; Posavi, M.; Hull, N.; Dinwiddie, D.; Halasz, A. M.; McGall, G.; Zhou, W.; Edwards, J. S. Highly Accurate Chip-Based Resequencing of SARS-CoV-2 Clinical Samples. Langmuir 2021. https://doi.org/10.1021/acs.langmuir.0c02927.

12. Quick, J. NCoV-2019 Sequencing Protocol v3 (LoCost). 2020.

13. Mockler, T. C.; Ecker, J. R. Applications of DNA Tiling Arrays for Whole-Genome Analysis. Genomics 2005, 85 (1), 1–15. https://doi.org/10.1016/j.ygeno.2004.10.005.

14. Shu, Y.; McCauley, J. GISAID: Global Initiative on Sharing All Influenza Data – from Vision to Reality. Euro Surveill 2017, 22 (13). https://doi.org/10.2807/1560-7917.ES.2017.22.13.30494.

15. Lanfear, R. A global phylogeny of SARS-CoV-2 sequences from GISAID. Zenodo DOI: 10.5281/zenodo.3958883. 2020.

16. H. Wickham. ggplot2: Elegant Graphics for Data Analysis. Springer-Verlag New York 2016

17. Chen, H.; Boutros, P. C. VennDiagram: A Package for the Generation of Highly-Customizable Venn and Euler Diagrams in R. BMC Bioinformatics 2011, 12 (1), 35. https://doi.org/10.1186/1471-2105-12-35.

18. Nowak, M. A. What Is a Quasispecies? Trends in Ecology & Evolution 1992, 7 (4), 118–121. https://doi.org/10.1016/0169-5347(92)90145-2.

19. Jary, A.; Leducq, V.; Malet, I.; Marot, S.; Klement-Frutos, E.; Teyssou, E.; Soulié, C.; Abdi, B.; Wirden, M.; Pourcher, V.; Caumes, E.; Calvez, V.; Burrel, S.; Marcelin, A.-G.; Boutolleau, D. Evolution of Viral Quasispecies during SARS-CoV-2 Infection. Clinical Microbiology and Infection 2020, 26 (11), 1560.e1–1560.e4. https://doi.org/10.1016/j.cmi.2020.07.032.

