## Supplemental Figures and Legends for all Clinical Samples for "SARS-CoV-2 Variant Identification Using a Genome Tiling Array and Genotyping Probes"

**Contents**
Number of pages: 26

Number of Tables: 1

Number of figures: 16

**Table S1:** Acknowledgement of laboratory/contributors of GISAID SARS-CoV-2 data

**Figure S1-S8**: Scatter plot of Q score and MAQ with variant call breakdown post MA-filter for each sample

**Figure S9-S16**: Venn diagram variant calls breakdown after MA-filter for each sample

**We gratefully acknowledge the following Authors from the Originating laboratories responsible for obtaining the specimens and the Submitting laboratories where genetic sequence data were generated and shared via the GISAID Initiative, on which this research is based.**

| **Virus Name** | **Accession No.** | **Collected** | **Originating Laboratory** | **Submitting laboratory** | **Authors** |
| --- | --- | --- | --- | --- | --- |
| hCoV-19/Wuhan/Hu-1/2019 | EPI_ISL_402125 | 12/31/2019 | National Institute for Communicable Disease Control and Prevention (ICDC) Chinese Center for Disease Control and Prevention (China CDC) | Chinese Center for Disease Control and Prevention | Zhang,Y.-Z., Wu,F., Chen,Y.-M., Pei,Y.-Y., Xu,L., Wang,W., Zhao,S., Yu,B., Hu,Y., Tao,Z.-W., Song,Z.-G., Tian,J.-H., Zhang,Y.-L., Liu,Y., Zheng,J.-J., Dai,F.-H., Wang,Q.-M., She,J.-L. and Zhu,T.-Y. |
| hCoV-19/USA/WY-WYPHL-00024/2020 | EPI_ISL_462917 | 3/22/2020 | Wyoming Public Health Laboratory | Center for Global Health, University of New Mexico Health Sciences Center | Daryl Domman, Kurt Schwalm, Rob Christensen, Wanda Manley, Cari Sloma, Noah Hull, Darrell Dinwiddie |
| hCoV-19/USA/WY-WYPHL-00026/2020 | EPI_ISL_462919 | 3/21/2020 | Wyoming Public Health Laboratory | Center for Global Health, University of New Mexico Health Sciences Center | Daryl Domman, Kurt Schwalm, Rob Christensen, Wanda Manley, Cari Sloma, Noah Hull, Darrell Dinwiddie |
| hCoV-19/USA/WY-WYPHL-00032/2020 | EPI_ISL_462925 | 3/23/2020 | Wyoming Public Health Laboratory | Center for Global Health, University of New Mexico Health Sciences Center | Daryl Domman, Kurt Schwalm, Rob Christensen, Wanda Manley, Cari Sloma, Noah Hull, Darrell Dinwiddie |
| hCoV-19/USA/CO-WYPHL-00036/2020 | EPI_ISL_462912 | 3/24/2020 | Wyoming Public Health Laboratory | Center for Global Health, University of New Mexico Health Sciences Center | Daryl Domman, Kurt Schwalm, Rob Christensen, Wanda Manley, Cari Sloma, Noah Hull, Darrell Dinwiddie |
| hCoV-19/USA/WY-WYPHL-00041/2020 | EPI_ISL_462933 | 3/24/2020 | Wyoming Public Health Laboratory | Center for Global Health, University of New Mexico Health Sciences Center | Daryl Domman, Kurt Schwalm, Rob Christensen, Wanda Manley, Cari Sloma, Noah Hull, Darrell Dinwiddie |
| hCoV-19/USA/WY-WYPHL-00044/2020 | EPI_ISL_462936 | 3/25/2020 | Wyoming Public Health Laboratory | Center for Global Health, University of New Mexico Health Sciences Center | Daryl Domman, Kurt Schwalm, Rob Christensen, Wanda Manley, Cari Sloma, Noah Hull, Darrell Dinwiddie |
| hCoV-19/USA/WY-WYPHL-00059/2020 | EPI_ISL_462951 | 3/26/2020 | Wyoming Public Health Laboratory | Center for Global Health, University of New Mexico Health Sciences Center | Daryl Domman, Kurt Schwalm, Rob Christensen, Wanda Manley, Cari Sloma, Noah Hull, Darrell Dinwiddie |
| hCoV-19/USA/WY-WYPHL-00064/2020 | EPI_ISL_462956 | 3/25/2020 | Wyoming Public Health Laboratory | Center for Global Health, University of New Mexico Health Sciences Center | Daryl Domman, Kurt Schwalm, Rob Christensen, Wanda Manley, Cari Sloma, Noah Hull, Darrell Dinwiddie |

**Table S1.** Acknowledgement and citation of SARS-CoV-2 sequencing data shared and found through GISAID Initiative.

**All submitters of data may be contacted directly via www.gisaid.org**

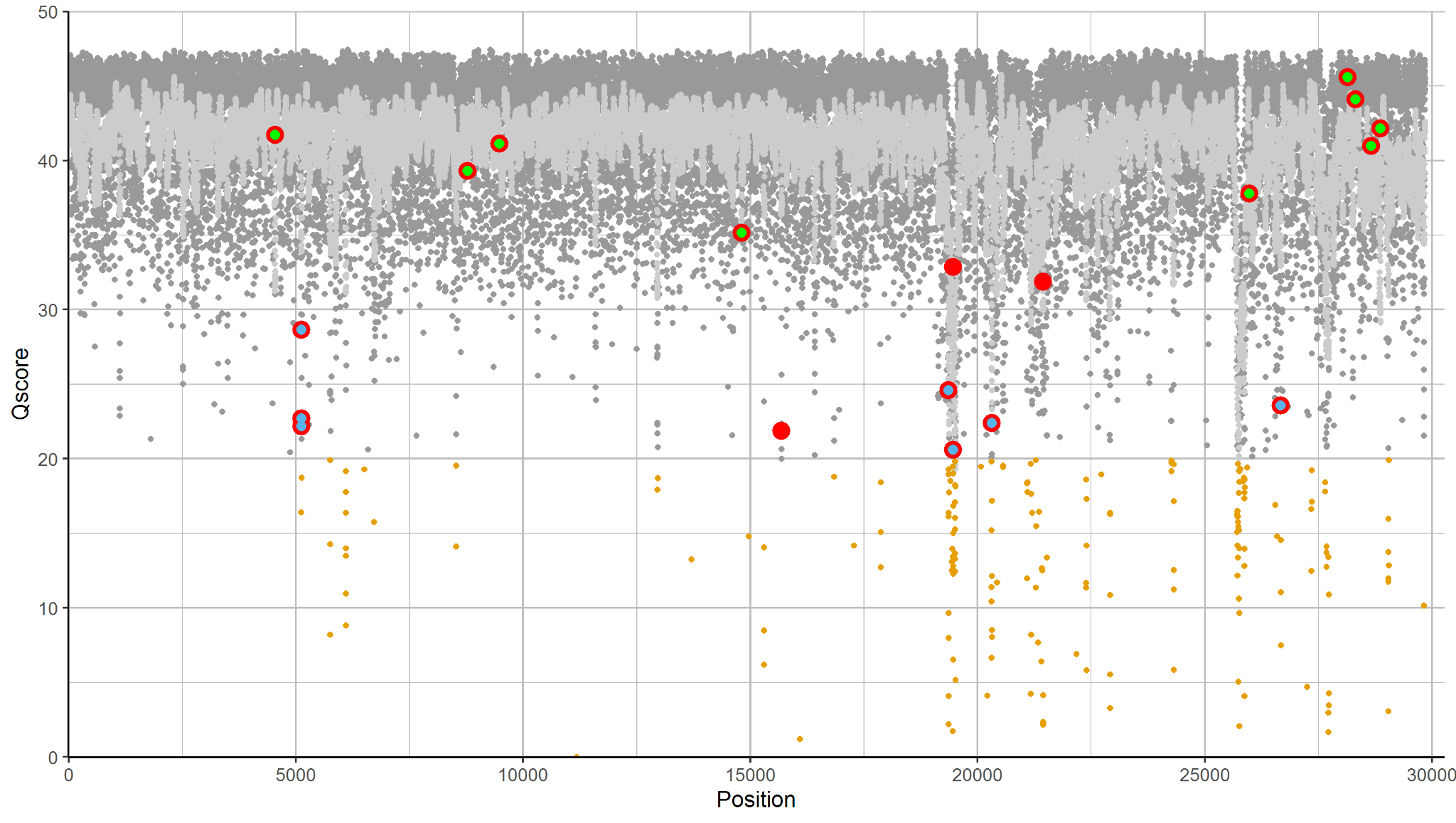

**Figure S1. Scatter plot, WY24 Q score and MAQ with variant call breakdown after MA-filter.**

A scatter plot generated in R using ggplot2 displaying all reads made by the full genome tiling array reads with incorporation of genotyping probe-set data and MA-filter on sample WY24 between position 26 and 29834. Every call was assessed with a combination of base call and Q score. High quality reads with Q > Qth (20) are used to make calls. Dark gray circles represent base positions where Q > Qth and make a ‘Reference Call’. All calculated MAQ are overlaid as light gray.  Brown dots are reads that have a low Q score where Q < Qth, are categorically ‘Non-calls’, and excluded to make base-calls. Q scores of ‘Variant Calls’ are identified as larger red circles, where the final base call made by DNA chip after replacement and MA-filter is not reference and the Q > Qth. Within ‘Variant Calls’, a blue overlap indicates calls that are removed by the MA-filter, which takes all variant calls with a Q score between 20 and 30 and removes any with a MAQ lower than the MA-threshold. Within ‘Variant Calls’, a green overlap indicates true variant calls identified and verified with short-read sequencing data.

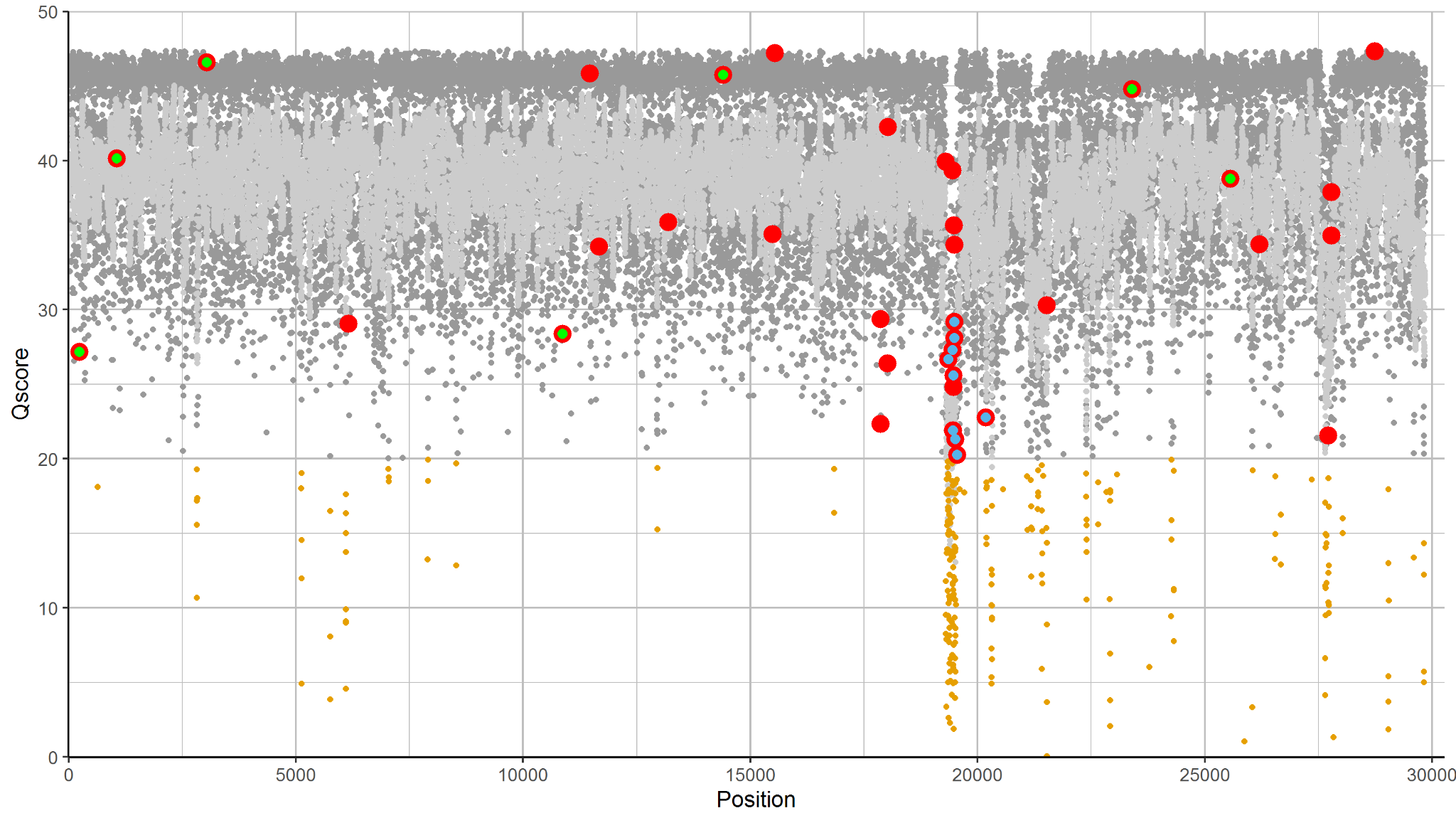

**Figure S2. Scatter plot, WY26 Q score and MAQ with variant call breakdown after MA-filter.**

A scatter plot generated in R using ggplot2 displaying all reads made by the full genome tiling array reads with incorporation of genotyping probe-set data and MA-filter on sample WY26 between position (X axis) 26 and 29834. Every call was assessed with a combination of base call and Q score. High quality reads with Q > Qth (20) are used to make calls. Dark gray circles represent base positions where Q > Qth and make a ‘Reference Call’. All calculated MAQ are overlaid as light gray.  Brown dots are reads that have a low Q score where Q < Qth, are categorically ‘Non-calls’, and excluded to make base-calls. Q scores of ‘Variant Calls’ are identified as larger red circles, where the final base call made by DNA chip after replacement and MA-filter is not reference and the Q > Qth. Within ‘Variant Calls’, a blue overlap indicates calls that are removed by the MA-filter, which takes all variant calls with a Q score between 20 and 30 and removes any with a MAQ lower than the MA-threshold. Within ‘Variant Calls’, a green overlap indicates true variant calls identified and verified with short-read sequencing data.

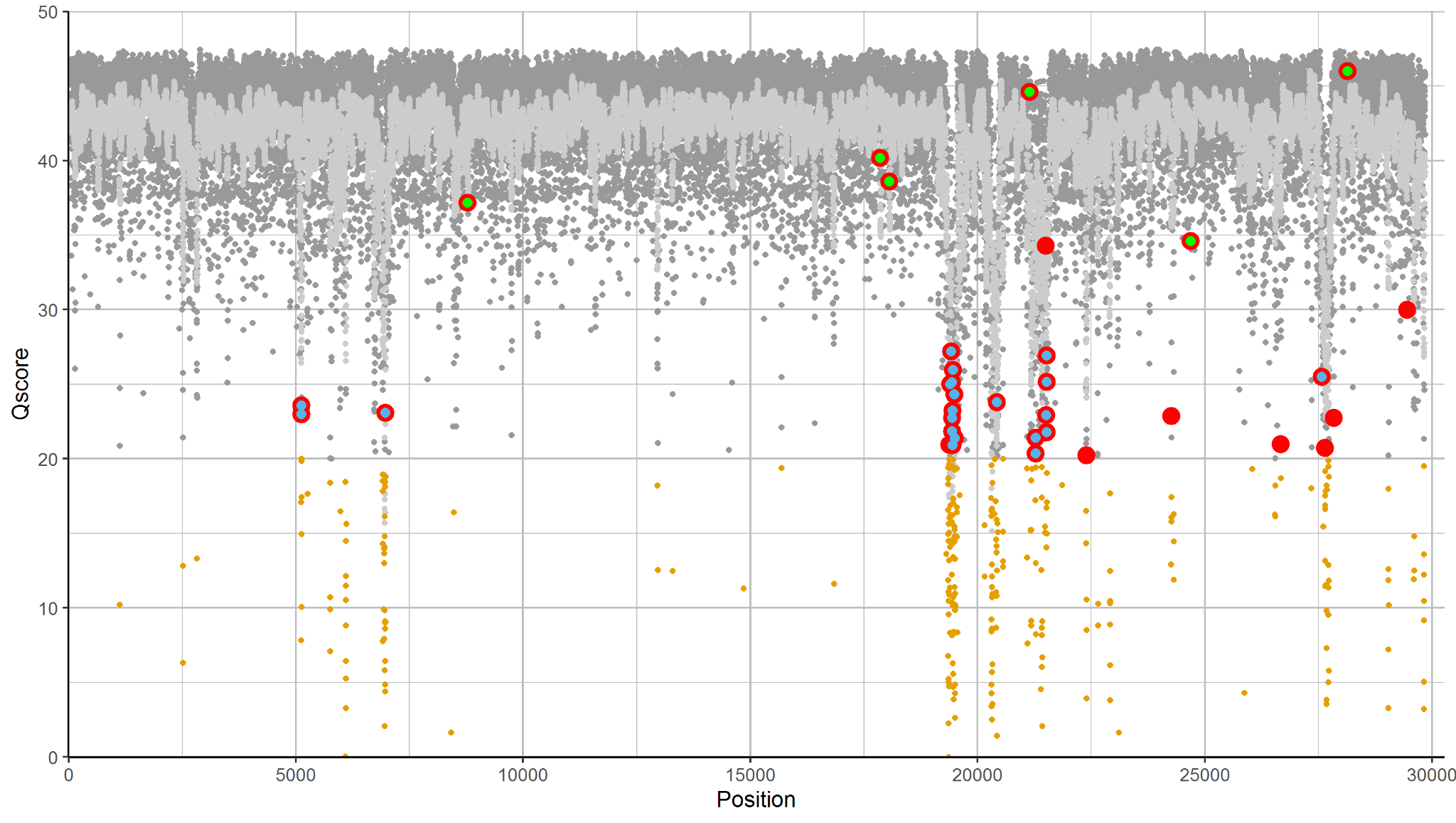

**Figure S3. Scatter plot, WY32 Q score and MAQ with variant call breakdown after MA-filter.**

A scatter plot generated in R using ggplot2 displaying all reads made by the full genome tiling array reads with incorporation of genotyping probe-set data and MA-filter on sample WY32 between position (X axis) 26 and 29834. Every call was assessed with a combination of base call and Q score. High quality reads with Q > Qth (20) are used to make calls. Dark gray circles represent base positions where Q > Qth and make a ‘Reference Call’. All calculated MAQ are overlaid as light gray.  Brown dots are reads that have a low Q score where Q < Qth, are categorically ‘Non-calls’, and excluded to make base-calls. Q scores of ‘Variant Calls’ are identified as larger red circles, where the final base call made by DNA chip after replacement and MA-filter is not reference and the Q > Qth. Within ‘Variant Calls’, a blue overlap indicates calls that are removed by the MA-filter, which takes all variant calls with a Q score between 20 and 30 and removes any with a MAQ lower than the MA-threshold. Within ‘Variant Calls’, a green overlap indicates true variant calls identified and verified with short-read sequencing data.

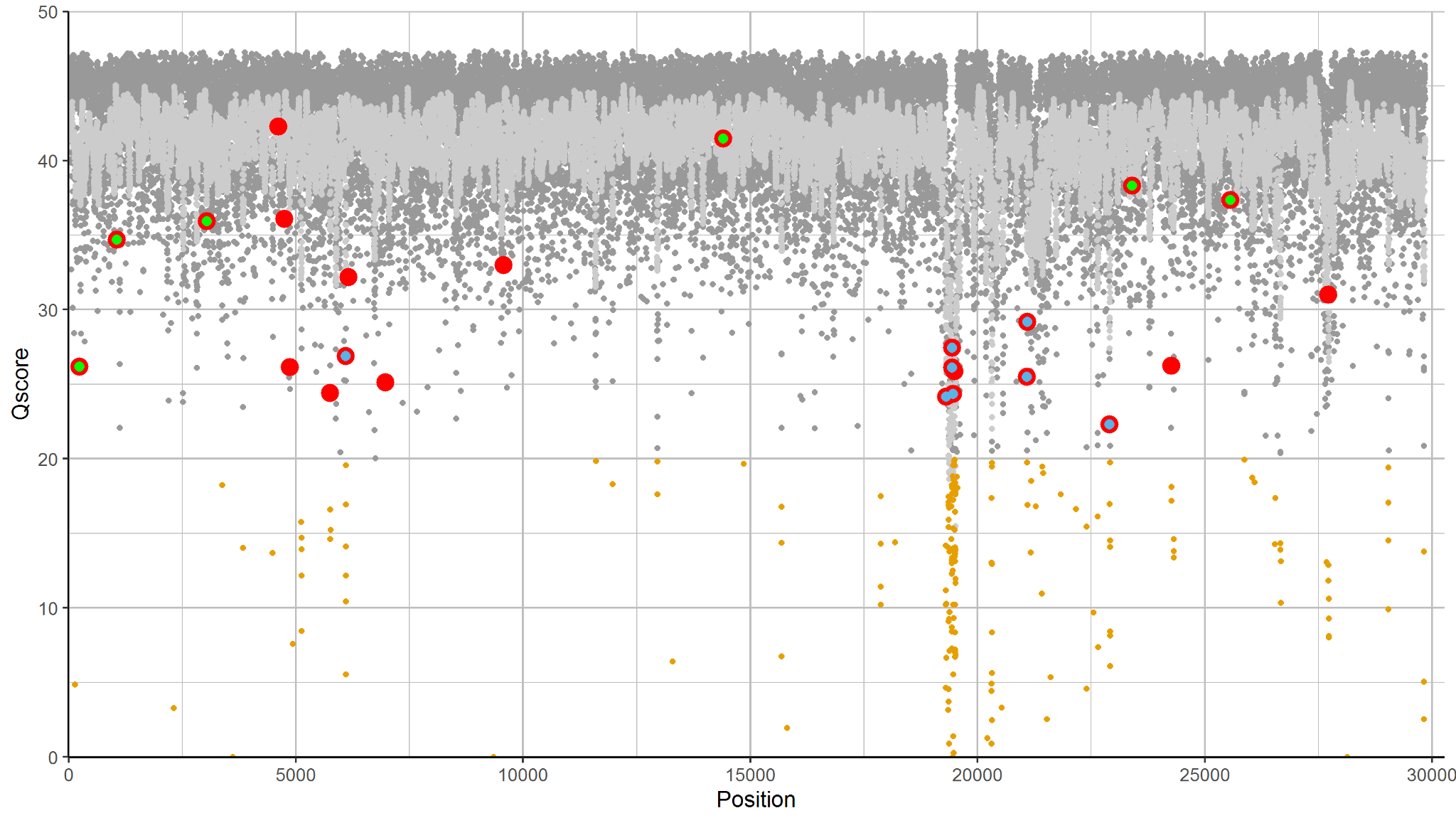

**Figure S4. Scatter plot, WY36 Q score and MAQ with variant call breakdown after MA-filter.**

A scatter plot generated in R using ggplot2 displaying all reads made by the full genome tiling array reads with incorporation of genotyping probe-set data and MA-filter on sample WY36 between position (X axis) 26 and 29834. Every call was assessed with a combination of base call and Q score. High quality reads with Q > Qth (20) are used to make calls. Dark gray circles represent base positions where Q > Qth and make a ‘Reference Call’. All calculated MAQ are overlaid as light gray.  Brown dots are reads that have a low Q score where Q < Qth, are categorically ‘Non-calls’, and excluded to make base-calls. Q scores of ‘Variant Calls’ are identified as larger red circles, where the final base call made by DNA chip after replacement and MA-filter is not reference and the Q > Qth. Within ‘Variant Calls’, a blue overlap indicates calls that are removed by the MA-filter, which takes all variant calls with a Q score between 20 and 30 and removes any with a MAQ lower than the MA-threshold. Within ‘Variant Calls’, a green overlap indicates true variant calls identified and verified with short-read sequencing data.

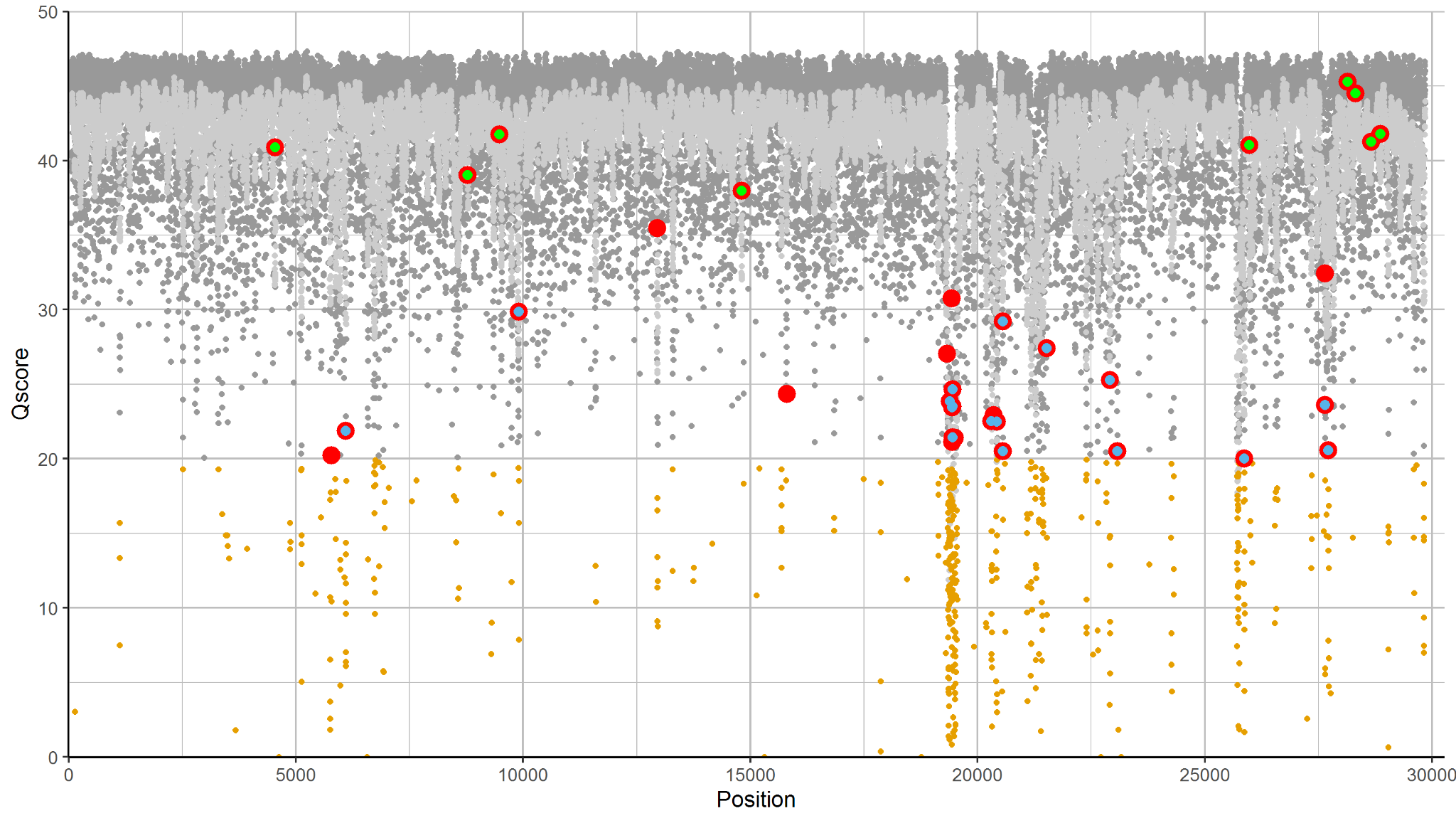

**Figure S5. Scatter plot, WY41 Q score and MAQ with variant call breakdown after MA-filter.**

A scatter plot generated in R using ggplot2 displaying all reads made by the full genome tiling array reads with incorporation of genotyping probe-set data and MA-filter on sample WY41 between position (X axis) 26 and 29834. Every call was assessed with a combination of base call and Q score. High quality reads with Q > Qth (20) are used to make calls. Dark gray circles represent base positions where Q > Qth and make a ‘Reference Call’. All calculated MAQ are overlaid as light gray.  Brown dots are reads that have a low Q score where Q < Qth, are categorically ‘Non-calls’, and excluded to make base-calls. Q scores of ‘Variant Calls’ are identified as larger red circles, where the final base call made by DNA chip after replacement and MA-filter is not reference and the Q > Qth. Within ‘Variant Calls’, a blue overlap indicates calls that are removed by the MA-filter, which takes all variant calls with a Q score between 20 and 30 and removes any with a MAQ lower than the MA-threshold. Within ‘Variant Calls’, a green overlap indicates true variant calls identified and verified with short-read sequencing data.

.

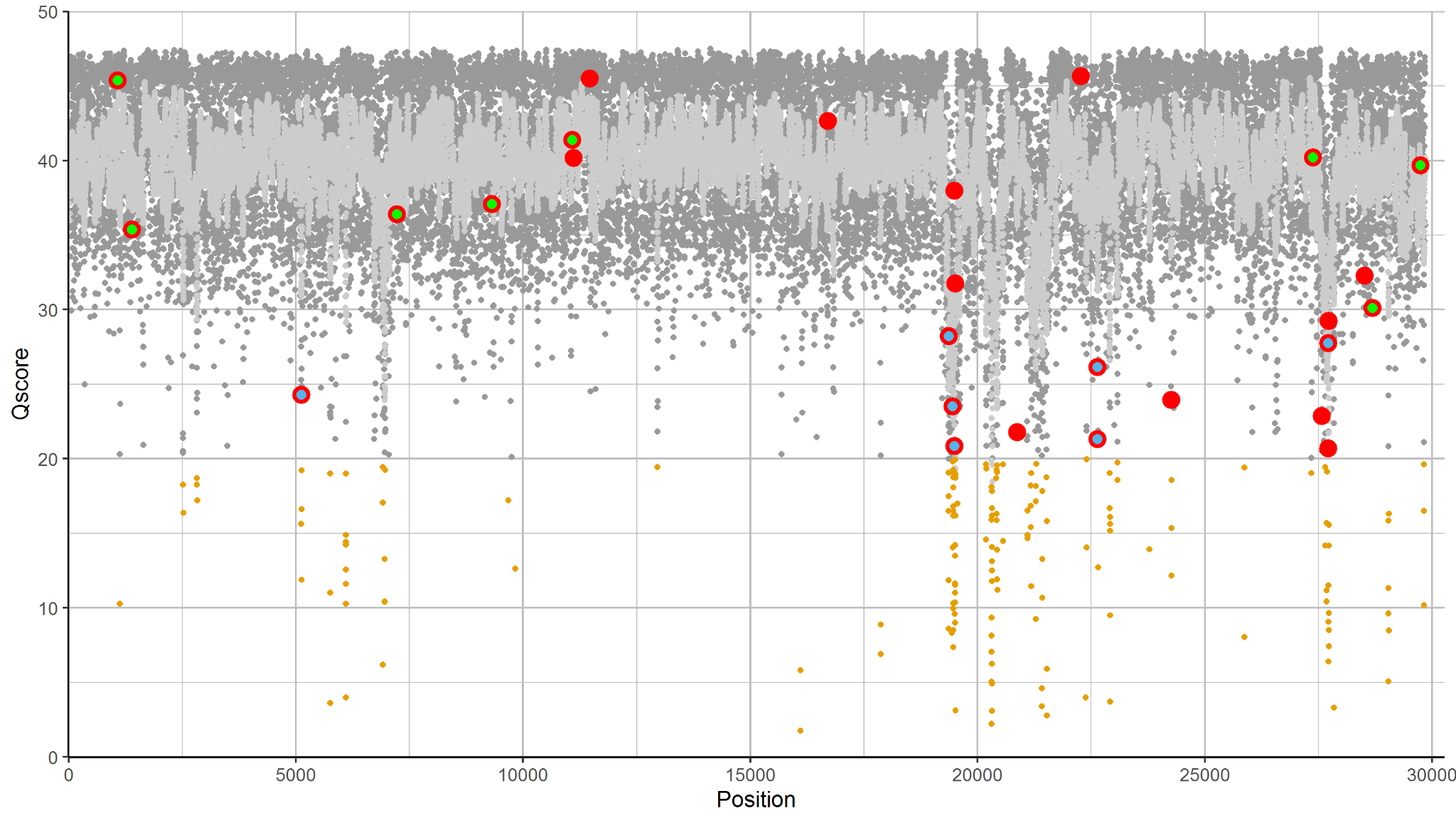

**Figure S6. Scatter plot, WY44 Q score and MAQ with variant call breakdown after MA-filter.**

A scatter plot generated in R using ggplot2 displaying all reads made by the full genome tiling array reads with incorporation of genotyping probe-set data and MA-filter on sample WY44 between position (X axis) 26 and 29834. Every call was assessed with a combination of base call and Q score. High quality reads with Q > Qth (20) are used to make calls. Dark gray circles represent base positions where Q > Qth and make a ‘Reference Call’. All calculated MAQ are overlaid as light gray.  Brown dots are reads that have a low Q score where Q < Qth, are categorically ‘Non-calls’, and excluded to make base-calls. Q scores of ‘Variant Calls’ are identified as larger red circles, where the final base call made by DNA chip after replacement and MA-filter is not reference and the Q > Qth. Within ‘Variant Calls’, a blue overlap indicates calls that are removed by the MA-filter, which takes all variant calls with a Q score between 20 and 30 and removes any with a MAQ lower than the MA-threshold. Within ‘Variant Calls’, a green overlap indicates true variant calls identified and verified with short-read sequencing data.

.
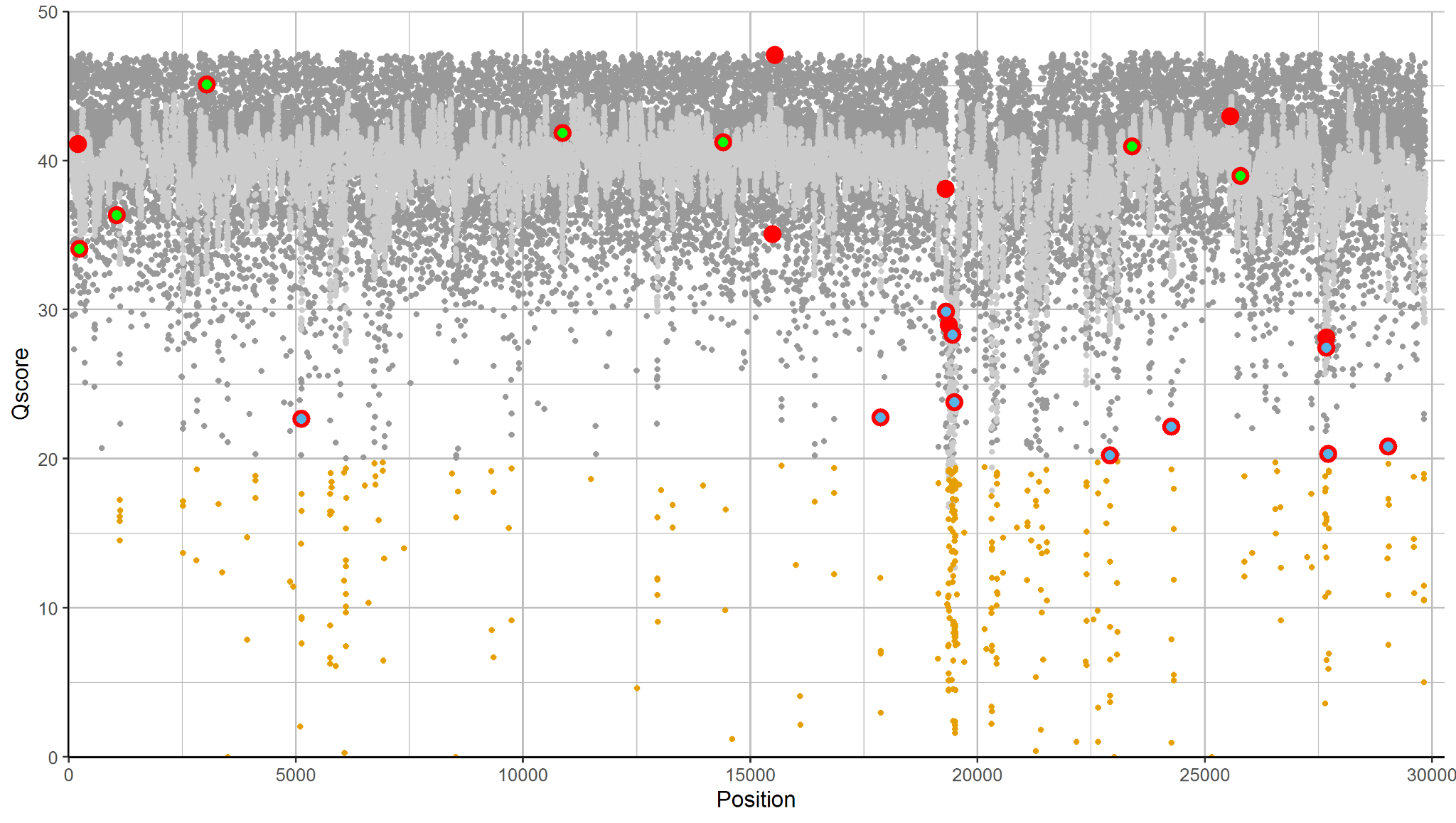

**Figure S7. Scatter plot, WY59 Q score and MAQ with variant call breakdown after MA-filter.**

A scatter plot generated in R using ggplot2 displaying all reads made by the full genome tiling array reads with incorporation of genotyping probe-set data and MA-filter on sample WY59 between position (X axis) 26 and 29834. Every call was assessed with a combination of base call and Q score. High quality reads with Q > Qth (20) are used to make calls. Dark gray circles represent base positions where Q > Qth and make a ‘Reference Call’. All calculated MAQ are overlaid as light gray.  Brown dots are reads that have a low Q score where Q < Qth, are categorically ‘Non-calls’, and excluded to make base-calls. Q scores of ‘Variant Calls’ are identified as larger red circles, where the final base call made by DNA chip after replacement and MA-filter is not reference and the Q > Qth. Within ‘Variant Calls’, a blue overlap indicates calls that are removed by the MA-filter, which takes all variant calls with a Q score between 20 and 30 and removes any with a MAQ lower than the MA-threshold. Within ‘Variant Calls’, a green overlap indicates true variant calls identified and verified with short-read sequencing data.

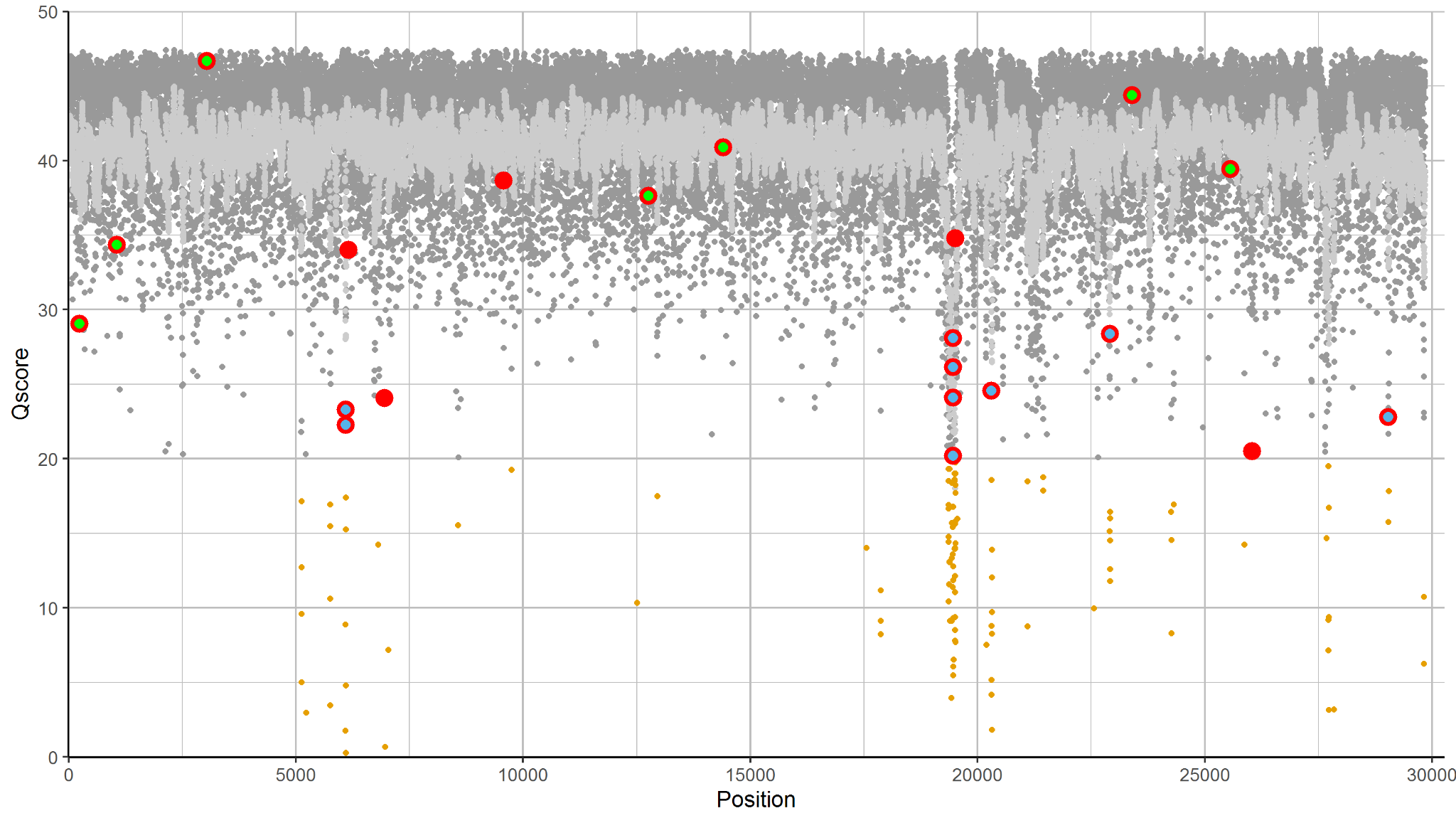

**Figure S8. Scatter plot, WY64 Q score and MAQ with variant call breakdown after MA-filter.**

scatter plot generated in R using ggplot2 displaying all reads made by the full genome tiling array reads with incorporation of genotyping probe-set data and MA-filter on sample WY64 between position (X axis) 26 and 29834. Every call was assessed with a combination of base call and Q score. High quality reads with Q > Qth (20) are used to make calls. Dark gray circles represent base positions where Q > Qth and make a ‘Reference Call’. All calculated MAQ are overlaid as light gray.  Brown dots are reads that have a low Q score where Q < Qth, are categorically ‘Non-calls’, and excluded to make base-calls. Q scores of ‘Variant Calls’ are identified as larger red circles, where the final base call made by DNA chip after replacement and MA-filter is not reference and the Q > Qth. Within ‘Variant Calls’, a blue overlap indicates calls that are removed by the MA-filter, which takes all variant calls with a Q score between 20 and 30 and removes any with a MAQ lower than the MA-threshold. Within ‘Variant Calls’, a green overlap indicates true variant calls identified and verified with short-read sequencing data.

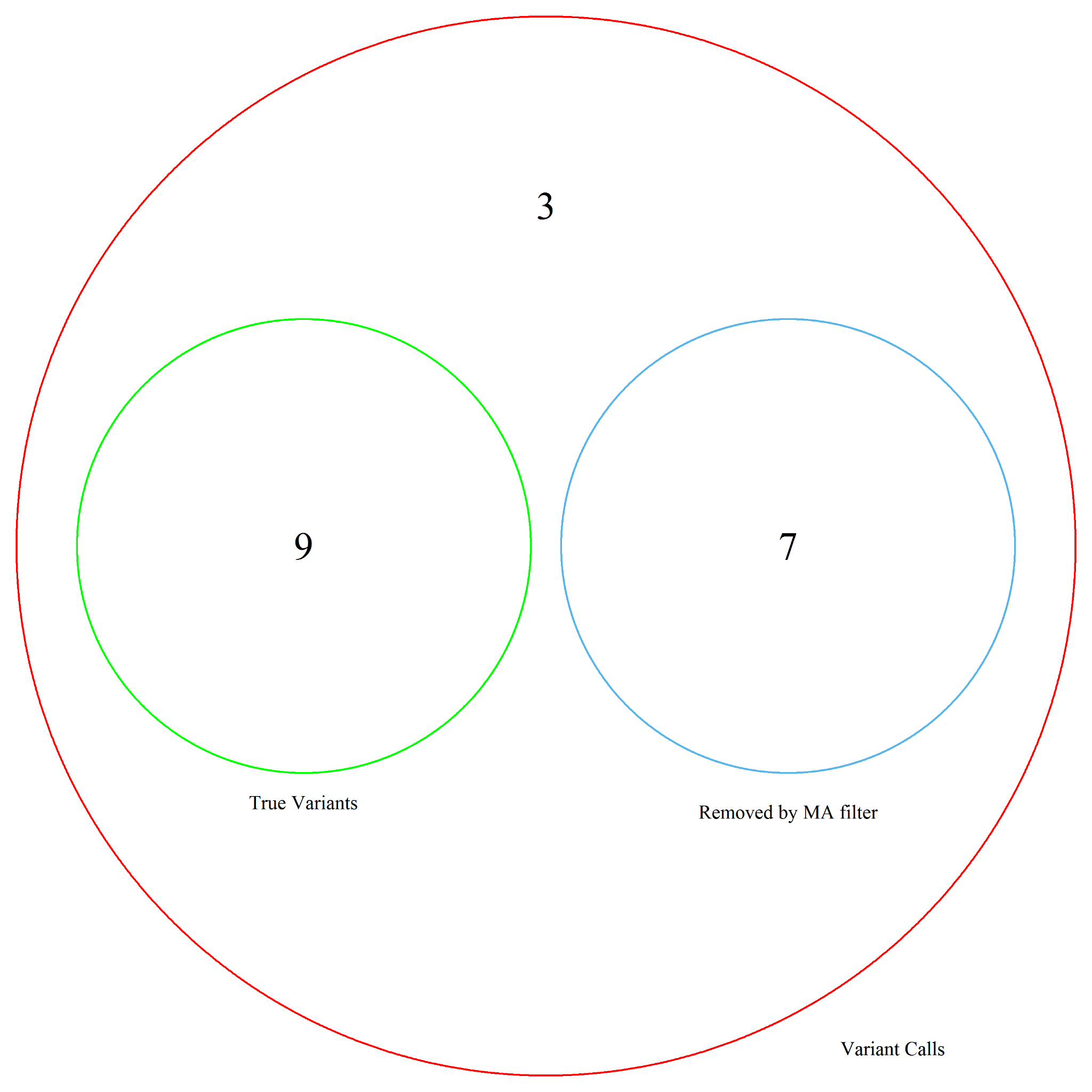
**Figure S9. Venn Diagram, WY24 Variant Calls after MA-filter.**

A Venn diagram depicting the categorical breakdown of calls made by the DNA chip on sample WY24. Variant Calls made by the DNA chip are contained within the red circle. Any circle or number outside of red implies that the call is a reference call. True variants are color coded in green and are confirmed by short-read sequencing data. The blue circle indicates variant calls that are filtered out by the MA-filter, converted to non-calls, and omitted from making base calls. Overlap between green and red circles indicates true variants correctly called by the DNA chip and verified through short-read sequencing data. Overlap between green and blue circles indicate an improperly removed true variant by the MA-filter where the read is situated in a local region of low MAs indicated by a low MAQ below the MAQ threshold. Ideally both green and blue circles should be contained within the red circle without overlapping each other, implying that all true variants and reads filtered out by the MA-filter were correctly identified as variant calls.

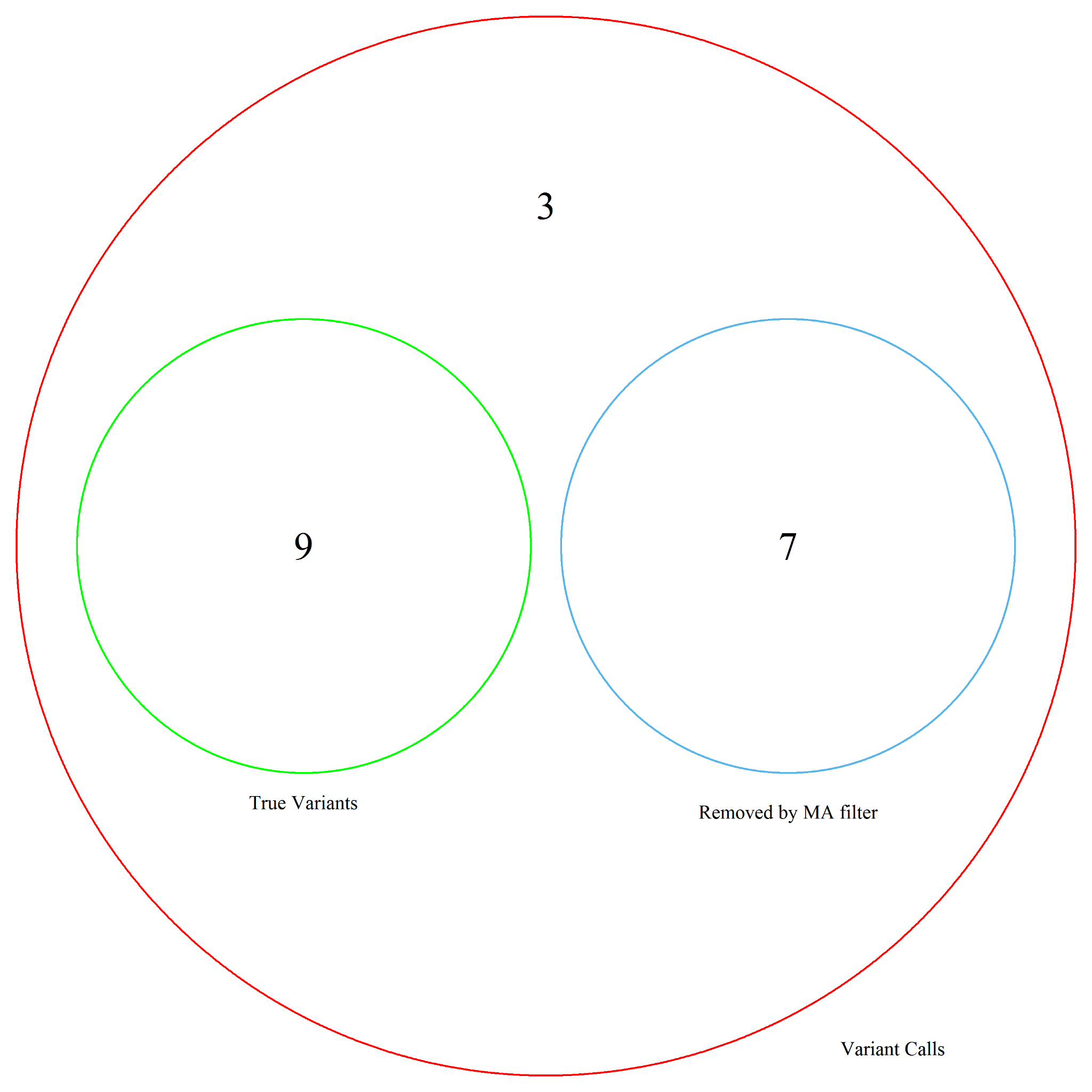
**Figure S10. Venn Diagram, WY26 Variant Calls after MA-filter.**

A Venn diagram depicting the categorical breakdown of calls made by the DNA chip on sample WY26. Variant Calls made by the DNA chip are contained within the red circle. Any circle or number outside of red implies that the call is a reference call. True variants are color coded in green and are confirmed by short-read sequencing data. The blue circle indicates variant calls that are filtered out by the MA-filter, converted to non-calls, and omitted from making base calls. Overlap between green and red circles indicates true variants correctly called by the DNA chip and verified through short-read sequencing data. Overlap between green and blue circles indicate an improperly removed true variant by the MA-filter where the read is situated in a local region of low MAs indicated by a low MAQ below the MAQ threshold. Ideally both green and blue circles should be contained within the red circle without overlapping each other, implying that all true variants and reads filtered out by the MA-filter were correctly identified as variant calls.

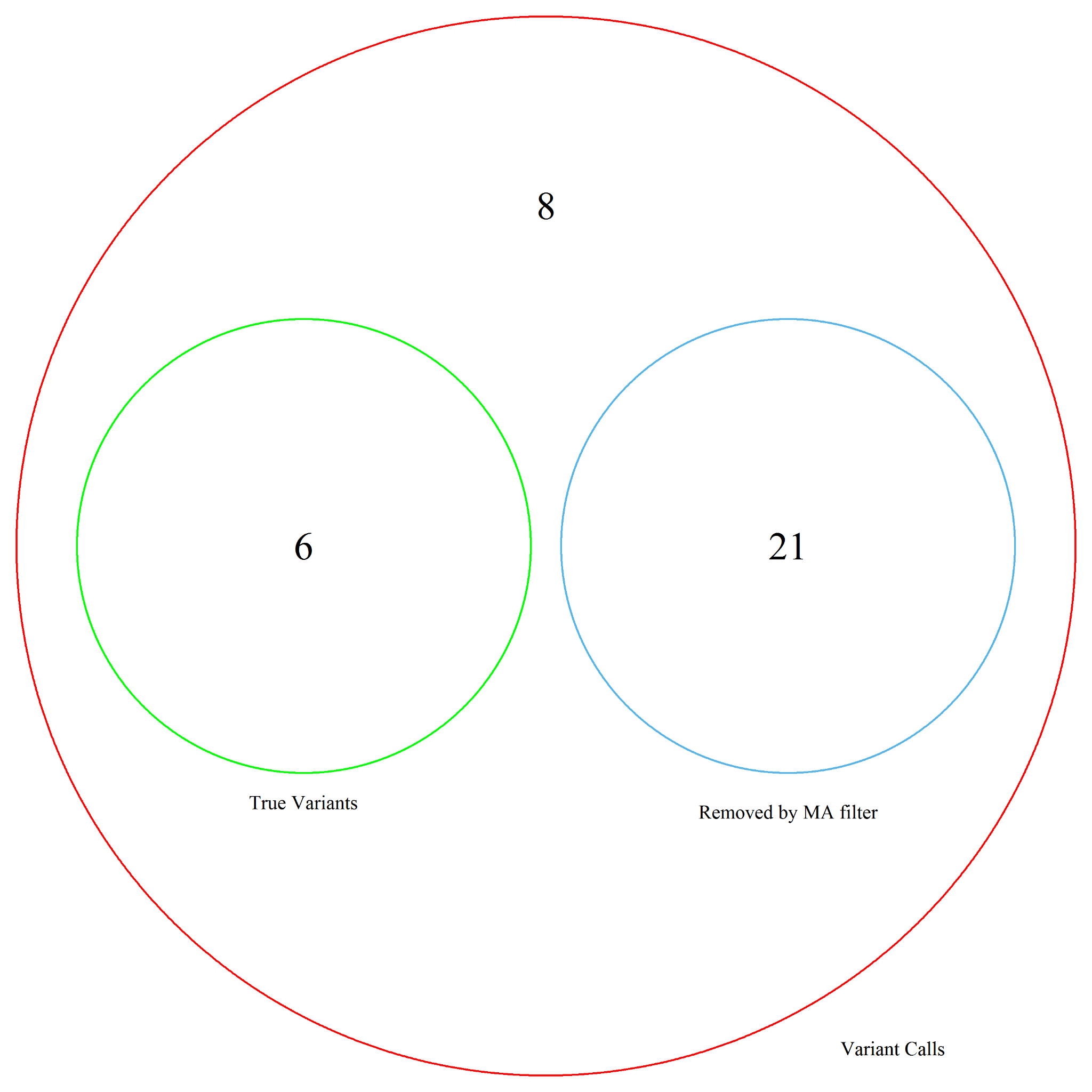
**Figure S11. Venn Diagram, WY32 Variant Calls after MA-filter.**

A venn diagram depicting the categorical breakdown of calls made by the DNA chip on sample WY32. Variant Calls made by the DNA chip are contained within the red circle. Any circle or number outside of red implies that the call is a reference call. True variants are color coded in green and are confirmed by short-read sequencing data. The blue circle indicates variant calls that are filtered out by the MA-filter, converted to non-calls, and omitted from making base calls. Overlap between green and red circles indicates true variants correctly called by the DNA chip and verified through short-read sequencing data. Overlap between green and blue circles indicate an improperly removed true variant by the MA-filter where the read is situated in a local region of low MAs indicated by a low MAQ below the MAQ threshold. Ideally both green and blue circles should be contained within the red circle without overlapping each other, implying that all true variants and reads filtered out by the MA-filter were correctly identified as variant calls.

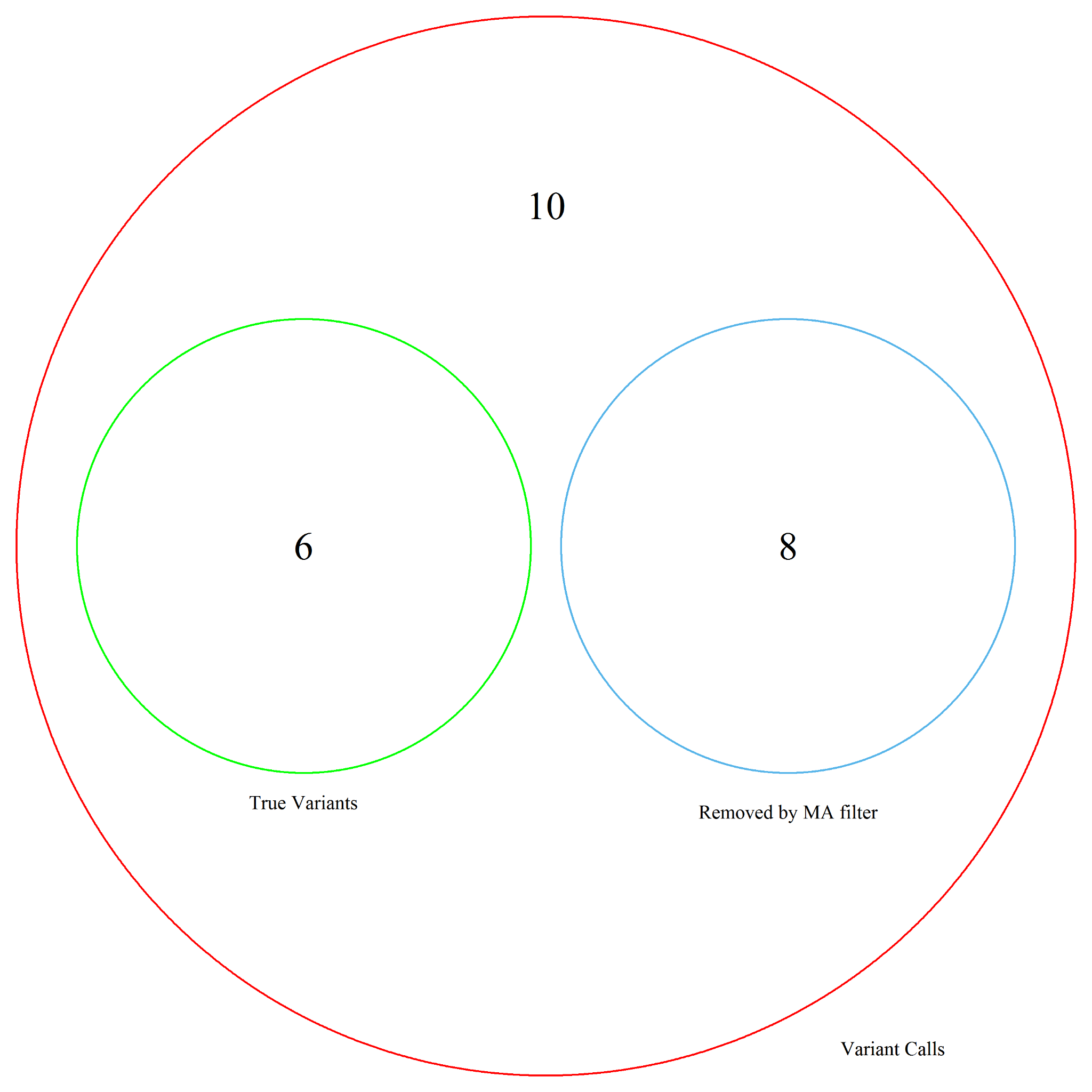
**Figure S12. Venn Diagram, WY36 Variant Calls after MA-filter.**

A venn diagram depicting the categorical breakdown of calls made by the DNA chip on sample WY36. Variant Calls made by the DNA chip are contained within the red circle. Any circle or number outside of red implies that the call is a reference call. True variants are color coded in green and are confirmed by short-read sequencing data. The blue circle indicates variant calls that are filtered out by the MA-filter, converted to non-calls, and omitted from making base calls. Overlap between green and red circles indicates true variants correctly called by the DNA chip and verified through short-read sequencing data. Overlap between green and blue circles indicate an improperly removed true variant by the MA-filter where the read is situated in a local region of low MAs indicated by a low MAQ below the MAQ threshold. Ideally both green and blue circles should be contained within the red circle without overlapping each other, implying that all true variants and reads filtered out by the MA-filter were correctly identified as variant calls.

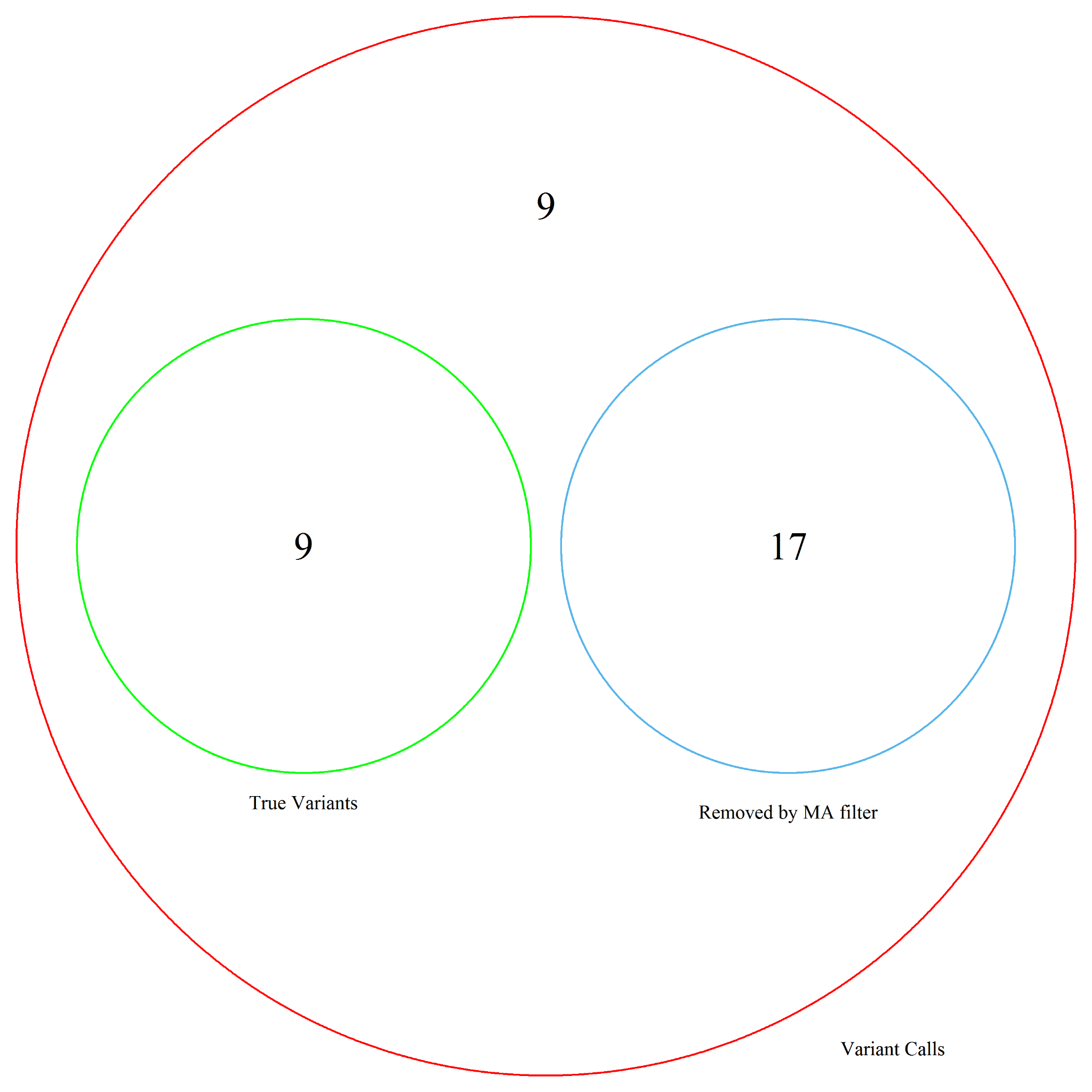
**Figure S13. Venn Diagram, WY41 Variant Calls after MA-filter.**

A venn diagram depicting the categorical breakdown of calls made by the DNA chip on sample WY41. Variant Calls made by the DNA chip are contained within the red circle. Any circle or number outside of red implies that the call is a reference call. True variants are color coded in green and are confirmed by short-read sequencing data. The blue circle indicates variant calls that are filtered out by the MA-filter, converted to non-calls, and omitted from making base calls. Overlap between green and red circles indicates true variants correctly called by the DNA chip and verified through short-read sequencing data. Overlap between green and blue circles indicate an improperly removed true variant by the MA-filter where the read is situated in a local region of low MAs indicated by a low MAQ below the MAQ threshold. Ideally both green and blue circles should be contained within the red circle without overlapping each other, implying that all true variants and reads filtered out by the MA-filter were correctly identified as variant calls.

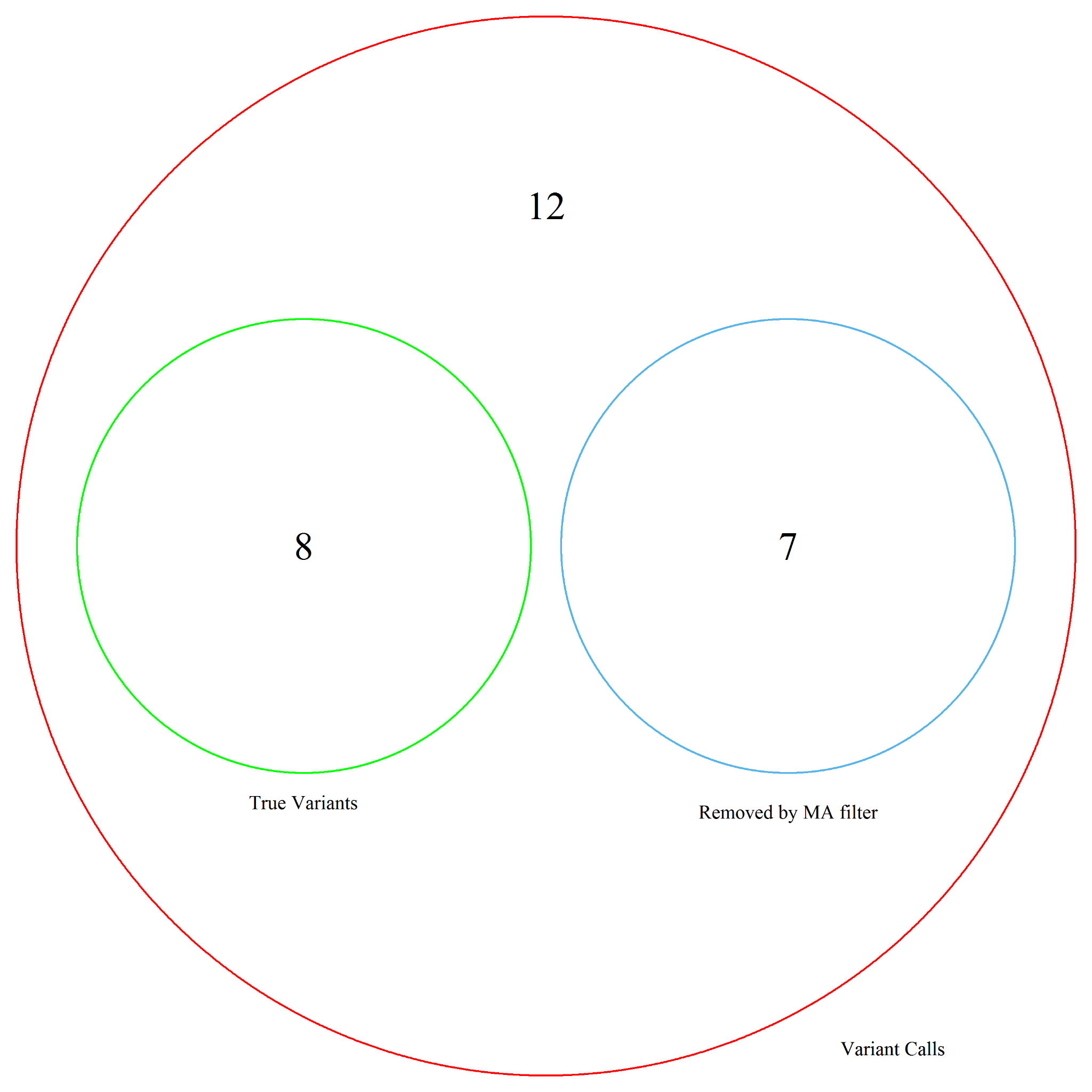

**Figures S14. Venn Diagram, WY44 Variant Calls after MA-filter.**

A venn diagram depicting the categorical breakdown of calls made by the DNA chip on sample WY44. Variant Calls made by the DNA chip are contained within the red circle. Any circle or number outside of red implies that the call is a reference call. True variants are color coded in green and are confirmed by short-read sequencing data. The blue circle indicates variant calls that are filtered out by the MA-filter, converted to non-calls, and omitted from making base calls. Overlap between green and red circles indicates true variants correctly called by the DNA chip and verified through short-read sequencing data. Overlap between green and blue circles indicate an improperly removed true variant by the MA-filter where the read is situated in a local region of low MAs indicated by a low MAQ below the MAQ threshold. Ideally both green and blue circles should be contained within the red circle without overlapping each other, implying that all true variants and reads filtered out by the MA-filter were correctly identified as variant calls.

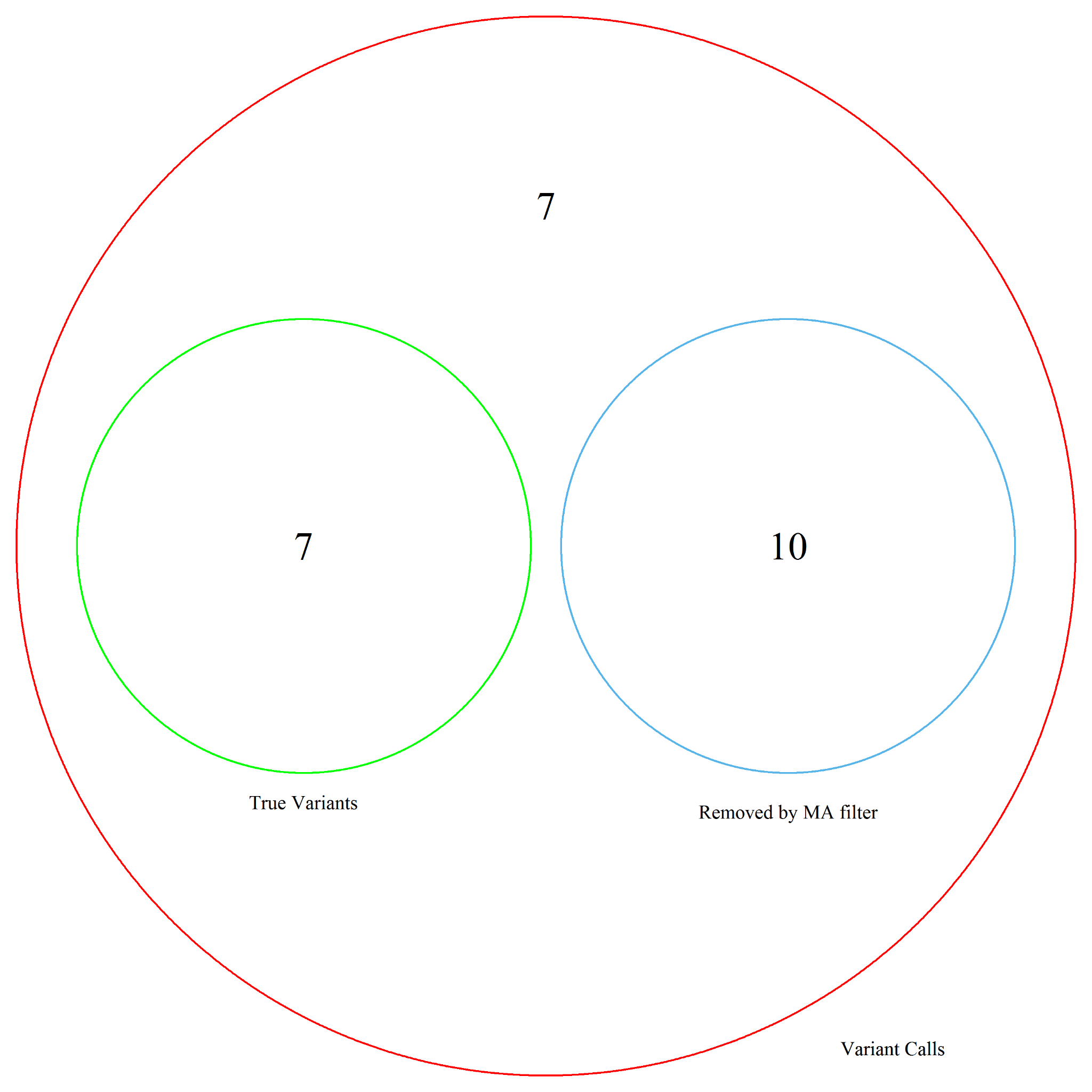
**Figure S15. Venn Diagram, WY59 Variant Calls after MA-filter.**

A venn diagram depicting the categorical breakdown of calls made by the DNA chip on sample WY59. Variant Calls made by the DNA chip are contained within the red circle. Any circle or number outside of red implies that the call is a reference call. True variants are color coded in green and are confirmed by short-read sequencing data. The blue circle indicates variant calls that are filtered out by the MA-filter, converted to non-calls, and omitted from making base calls. Overlap between green and red circles indicates true variants correctly called by the DNA chip and verified through short-read sequencing data. Overlap between green and blue circles indicate an improperly removed true variant by the MA-filter where the read is situated in a local region of low MAs indicated by a low MAQ below the MAQ threshold. Ideally both green and blue circles should be contained within the red circle without overlapping each other, implying that all true variants and reads filtered out by the MA-filter were correctly identified as variant calls.

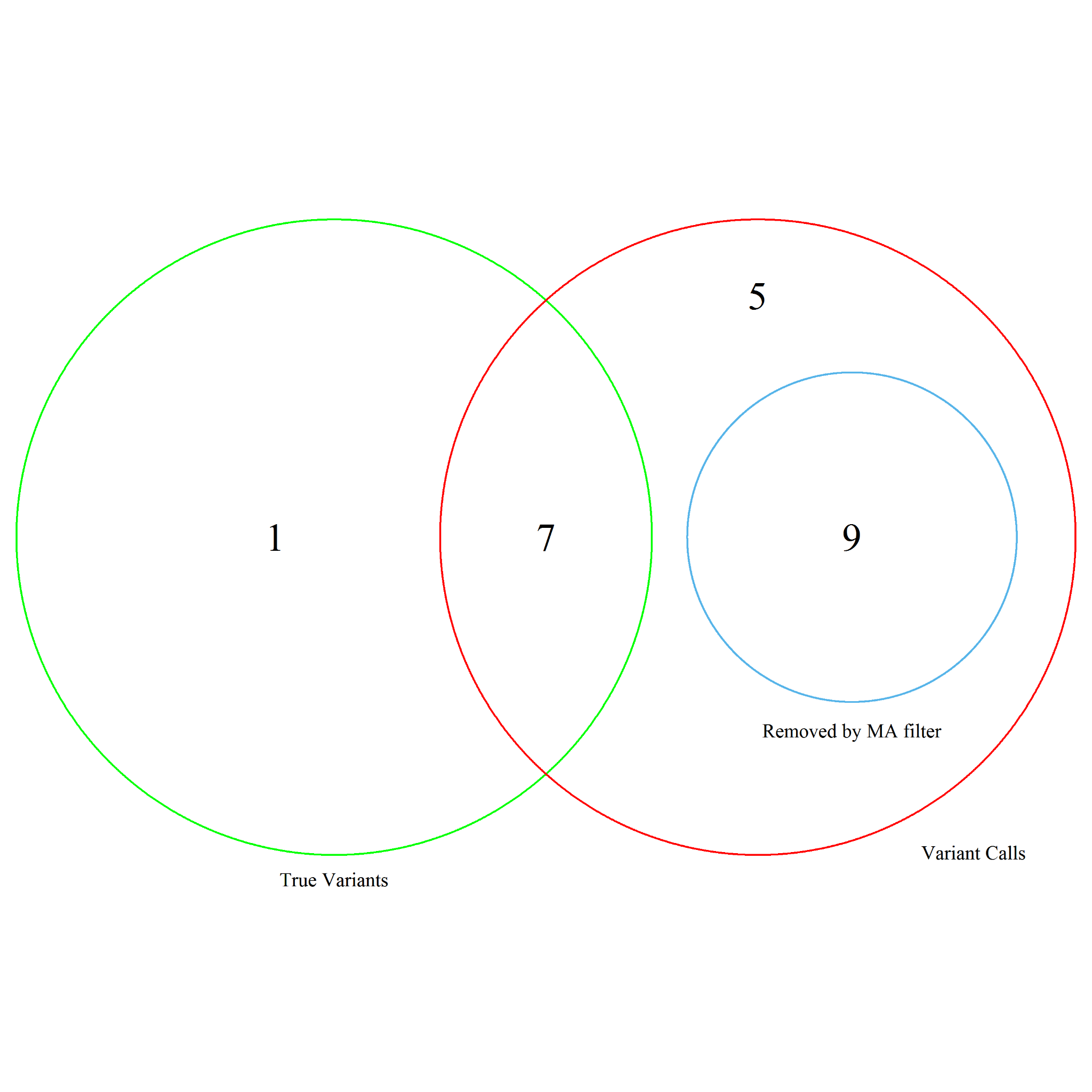
**Figure S16. Venn Diagram, WY64 Variant Calls after MA-filter.**

A venn diagram depicting the categorical breakdown of calls made by the DNA chip on sample WY64. Variant Calls made by the DNA chip are contained within the red circle. Any circle or number outside of red implies that the call is a reference call. True variants are color coded in green and are confirmed by short-read sequencing data. The blue circle indicates variant calls that are filtered out by the MA-filter, converted to non-calls, and omitted from making base calls. Overlap between green and red circles indicates true variants correctly called by the DNA chip and verified through short-read sequencing data. Overlap between green and blue circles indicate an improperly removed true variant by the MA-filter where the read is situated in a local region of low MAs indicated by a low MAQ below the MAQ threshold. Ideally both green and blue circles should be contained within the red circle without overlapping each other, implying that all true variants and reads filtered out by the MA-filter were correctly identified as variant calls.
